# Smoothened receptor Signaling regulates the developmental shift of GABA polarity in rat somatosensory cortex

**DOI:** 10.1101/799015

**Authors:** Quentin Delmotte, Igor Medina, Mira Hamze, Emmanuelle Buhler, Jinwei Zhang, Yesser H. Belgacem, Christophe Porcher

**Affiliations:** Aix-Marseille Univ, INSERM, INMED, Parc Scientifique de Luminy 13273, Marseille, France; INSERM (Institut National de la Santé et de la Recherche Médicale) Unité 1249, Marseille, Parc Scientifique de Luminy, 13273 Marseille, France; INMED (Institut de Neurobiologie de la Méditerranée), Parc Scientifique de Luminy, 13273 Marseille, France; Plateforme Post-Génomique, INMED, 13273 Marseille, France; Institute of Biomedical and Clinical Sciences, Medical School, College of Medicine and Health, University of Exeter, Hatherly Laboratories, Exeter, EX4 4PS, UK

**Keywords:** Smoothened receptor, Sonic hedgehog, KCC2, chloride homeostasis, GABAergic transmission

## Abstract

Sonic Hedgehog (Shh) and its patched-smoothened receptor complex control a variety of functions in the developing central nervous system (CNS) such as neural cell proliferation and differentiation. Recently, Shh signaling components have been found to be expressed at the synaptic level in the postnatal brain, suggesting a potential role in the regulation of synaptic transmission. Using *in utero* electroporation of constitutively active and dominant-negative forms of the Shh co-receptor smoothened (Smo), we studied the role of Smo signaling in the development and maturation of GABAergic transmission in the somatosensory cortex. Our results show that enhancing Smo activity during development accelerates the shift from depolarizing to hyperpolarizing GABA in dependence on functional expression of potassium-chloride cotransporter type 2 (KCC2). On the other hand, blocking Smo activity maintains GABA response in a depolarizing state in mature cortical neurons resulting in altered chloride homeostasis and increased seizure susceptibility. This study reveals an unexpected function of Smo signaling on the regulation of chloride homeostasis through the control of KCC2 cell surface stability and on the timing of the GABA inhibitory/excitatory shift in brain maturation.

**Summary statement:** The smoothened receptor controls the time course of inhibitory transmission through the stability of the potassium-chloride cotransporter type 2 at the plasma membrane.

## Introduction

Sonic hedgehog (Shh) is an activity-dependent secreted synaptic molecule (Beug et al., 2011). In the presence of Shh ligand, receptor Patched-1 (Ptch1) relieves its constitutive inhibition on G-protein coupled co-receptor Smoothened (Smo), leading to activation of downstream signaling factors (Belgacem and Borodinsky, 2015; Briscoe and Thérond, 2013; Riobo et al., 2006). Shh signaling plays a wide range of functions during embryonic development from neuronal cell proliferation to differentiation (Belgacem et al., 2016; Briscoe and Thérond, 2013; Ruat et al., 2012; Traiffort et al., 2010). Furthermore, in the developing mammalian central nervous system, Shh signaling is involved in axonal elongation (Hammond et al., 2009; Parra and Zou, 2010; Yao et al., 2015) and formation of the cortical connectivity (Harwell et al., 2012; Memi et al., 2018). After birth, Shh signaling components have been found in several brain regions, including the cerebral cortex and hippocampus, from an early stage of postnatal development to adulthood (Charytoniuk et al., 2002; Petralia et al., 2011; Petralia et al., 2012; Rivell et al., 2019; Traiffort et al., 2010). In the adult hippocampus, Shh plays an essential role in neurogenesis in the dentate gyrus (Antonelli et al., 2019; Breunig et al., 2008). In addition, Shh receptors Ptch1 and Smo are also expressed at the synaptic junction of the immature and adult hippocampus (Charytoniuk et al., 2002; Mitchell et al., 2012; Petralia et al., 2011), suggesting other roles than control of neurogenesis (Zuñiga and Stoeckli, 2017). For instance, Shh was reported to exert a modulatory action on neuronal electrical activity in the adult brain (Bezard et al., 2003; Pascual et al., 2005) and more recently Shh signaling has been shown to regulates the formation of glutamatergic and GABAergic terminals in hippocampal neurons (Mitchell et al., 2012). These morphological changes were accompanied by an increase in the frequency of excitatory postsynaptic currents (Feng et al., 2016; Mitchell et al., 2012). Altogether, these findings support the view that the Shh signaling pathway plays a crucial role during early postnatal neuronal circuit construction and synaptic plasticity in the hippocampus and cortex. Thus, its impairment may affect neuronal network formation and lead to brain dysfunction. Likewise, both in human and animal models accumulating evidences indicate that impairment of Shh pathway at postnatal stages may contribute for the emergence of neurodevelopmental disorders, including autism spectrum disorders (ASD) (Al-Ayadhi, 2012; Halepoto et al., 2015) and seizures (Feng et al., 2016; Su et al., 2017).

In the early postnatal life, the GABA_A_ receptor function changes from depolarizing to hyperpolarizing in a KCC2 dependent manner (Ben-Ari et al., 1989; Rivera et al., 1999). Alteration in the GABA developmental sequence leads to compromised balance between excitatory and inhibitory transmission associated with severe changes in the neuronal network circuitry (Ben-Ari, 2002; Ben-Ari et al., 2007; Kilb et al., 2013; Sernagor et al., 2010; Wang and Kriegstein, 2009) and subsequently, the onset of neurodevelopmental brain disorders (Ben-Ari and Holmes, 2005; Kuzirian and Paradis, 2011; Mueller et al., 2015).

Although both Shh and GABA signaling are critically involved in similar processes of neuronal network formation and are implicated in etiology of same neurodevelopmental disorders, their putative interplay remains unclear. Here we postulated that Shh components may contribute to the onset of the GABA polarity shift and suggested that deregulation of Shh signaling pathway may then lead to an impaired neuronal network and to the emergence of brain disorders at adulthood. Because loss of function of Smo have an embryonic lethal phenotype (Zhang et al., 2001) and Smo activation in Wnt1-Cre;R26SmoM2 mice induces a hyperplasia of the facial processes at embryonic stages (Jeong et al., 2004), we used *in utero* electroporation (IUE) to target the somatosensory cortex with constructs encoding either for the dominant-negative (Smo Δ570-581, called Smo-DN) (Kim et al., 2009) or the constitutively active (Smo A1, called Smo-CA) (Chen et al., 2011) Smo variants. Our results reveal an unexpected function for Shh/Ptch/Smo signaling pathway in the regulation of GABAergic developmental timing in the rat cerebral cortex, notably through the modulation of cell surface expression and stability of KCC2. We further show that impairment of Shh signaling alters the phosphorylated state of KCC2, thus maintaining a depolarizing action of GABA, which leads to an alteration of GABAergic inhibitory transmission and may contribute to the emergence of brain disorders.

## Results

### Cortical expression pattern of Smo-CA and Smo-DN

Previous studies have shown that Shh signaling pathways components, including Ptch1 and Smo, are present in the postnatal mammalian brain (Harwell et al., 2012; Mitchell et al., 2012; Petralia et al., 2011; Rivell et al., 2019). In the mouse neocortex, Shh signaling regulates the formation of neuronal network in immature brain, suggesting a potential function in construction and maturation of neural circuits during the postnatal periods. In order to evaluate its role in the developing brain, we used IUE of plasmids to express GFP alone as control or in combination with constitutively active (Smo-CA) or dominant-negative (Smo-DN) forms of Smo, an essential component for Shh signaling. IUE of rat brains was performed at embryonic day 15 (E15) in order to target mainly pyramidal neurons of the layers V/VI (Kriegstein and Noctor, 2004). Coronal sections through the somatosensory cortex of GFP and Smo-related constructs co-electroporated tissues from P15 rodents were immunolabeled with neuronal markers NeuN and FoxP2 (deeper layer neurons markers) to assist in the histological identification of cortical layers (Fig. 1A). The fluorescence intensity profile of both Smo-CA and Smo-DN expressing neurons does not show any obvious differences in the cortical layering when compared to control GFP-expressing cells. Thus, Smo-CA or Smo-DN expressing neurons electroporated at E15 most likely represent the layers V/VI of the somatosensory cortex.

**Figure 1:**
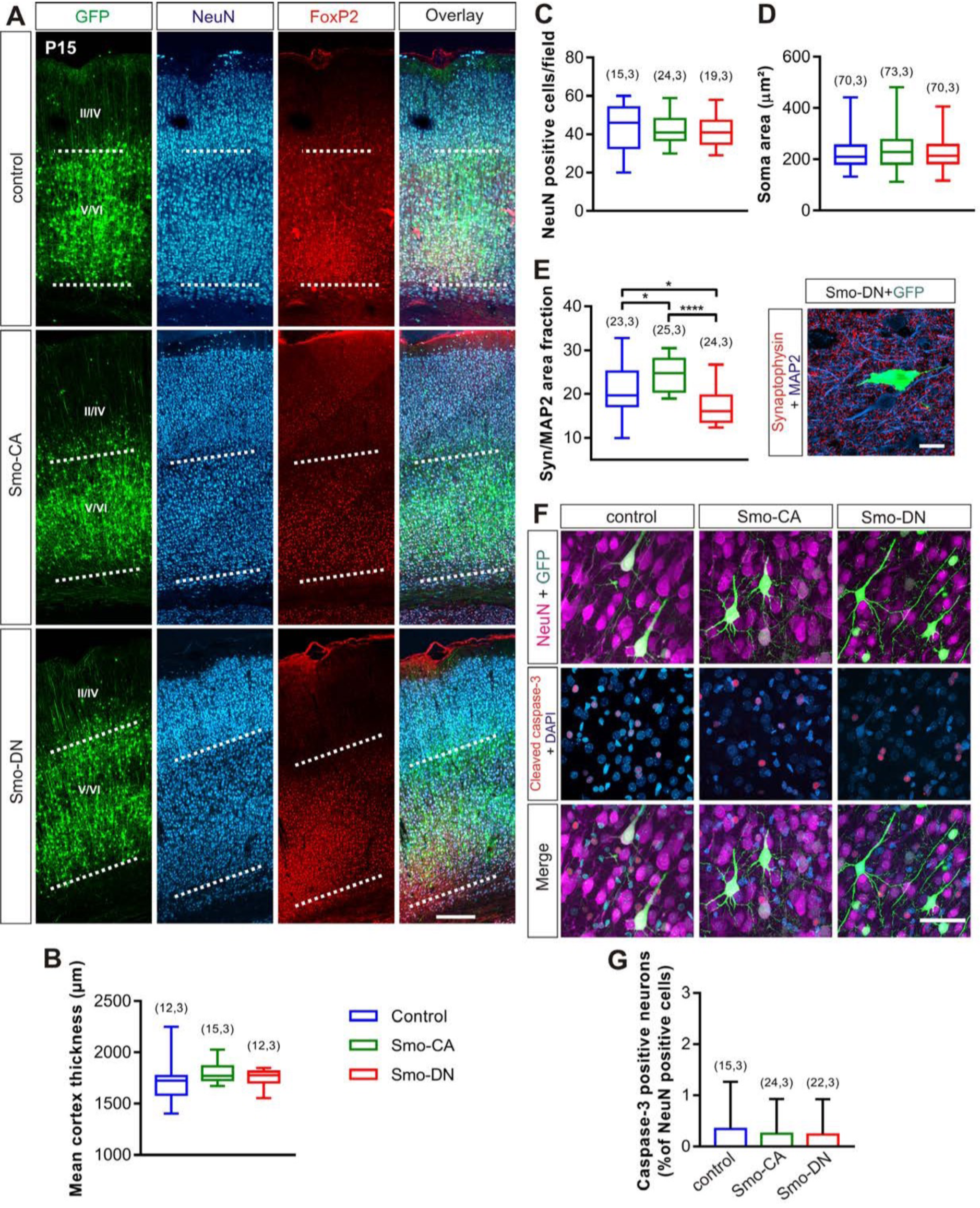
Cortical expression pattern of Smo-related constructs. (**A**) Representative neocortical section showing laminar position of transfected cells electroporated at E15 with control GFP (control) or co-electroporated with Smo-CA (Smo-CA) or Smo-DN (Smo-DN) and processed postnatally at P15 and co-immunostained with NeuN (blue) and FoxP2 (red). GFP fluorescence intensity profile shows that E15 generated neurons are distributed mainly in deeper layers V-VI. Scale bar = 250 µm. (**B**) Box plots of cortical thickness in GFP (control), Smo-CA and Smo-DN brains. Mann-Whitney test. (**C**) Box plots of neuronal density in GFP (control), Smo-CA and Smo-DN brains. (**D**) Box plots of neurons soma size on P15 electroporated rats. (**E**) Box plots of synaptophysin area fraction normalized with MAP2 area fraction (left) and (right) example of immunofluorescence signal of P30 cortical section expressing Smo-DN + GFP, co-immunostained with Synaptophysin (red) and MAP2 (blue). Scale bar = 30 µm. (**F**) Immunofluorescence signal of P15 cortical sections electroporated at E15 with GFP (control) with or without Smo-related constructs and immunolabeled with cleaved caspase-3 antibody. Scale bar = 100 µm. (**G**) Ratio of cleaved caspase-3-positive neurons (apoptotic) per surface area and expressed as a percentage of control (GFP). Overexpression of Smo-CA or Smo-DN does not increase the number of apoptotic neurons compared to control (GFP). Errors bars are mean ± SD. Number of slices and rats are indicated in parenthesis. \**p* < 0.05; \*\**p* < 0.01; \*\*\**p* < 0.001.

Because Shh signaling is involved in cell division and cell growth of cerebral cortex progenitors (Araújo et al., 2014), we investigated whether Smo-related constructs could impact the cortical thickness or the neuronal cell density and size in electroporated tissues. We found that cortical thickness of Smo-CA and Smo-DN rats does not present any difference compared to control rats (median values: 1772 µm, 1779 µm and 1772 µm respectively; *p* = 0.07 between Smo-CA and control rats, *p* = 0.17 between Smo-DN and control, Mann-Whitney test; Fig. 1B). The density of NeuN immuno-positive cells in Smo-CA and Smo-DN electroporated somatosensory cortices, measured in the layer V, reveals no difference when compared to control tissues (46 neurons per observation field for control, vs 41 neurons for Smo CA; *p* = 0.72 and 41 neurons for Smo-DN; *p* = 0.44, Mann-Whitney test; Fig. 1C). Similarly, the average soma neuronal size remains unchanged (210,3 µm^2^ for control vs 229.2 µm^2^ for Smo-CA and 213.8 µm^2^ for Smo-DN; *p* = 0.26 and *p* = 0.48 between control and Smo-DN, Mann-Whitney test; Fig. 1D).

In hippocampal neurons, exogenous treatment with Shh or Smo agonist (SAG) upregulate presynaptic nerve terminals (Mitchell et al., 2012). These observations led us to investigate whether Smo-CA or Smo-DN could modulate the density of presynaptic terminals in electroporated cortices. In order to assess the possible changes at the presynaptic levels, we performed quantitative analysis of the area fraction of synaptophysin on MAP2-positive neurons at P30. Compared with control GFP, Smo-DN electroporated tissues showed a significant decrease in the area fraction of synaptophysin immunolabeling (16.05 % for Smo-DN vs 19.67 % for control; *p* = 0.01, Mann-Whitney test; Fig. 1E), whereas Smo-CA increased the area fraction of synaptophysin-positive synapses (24.77 % for Smo-CA; *p* = 0.01 when compared to control and *p* < 0.0001 when compared to Smo-DN, Mann-Whitney test; Fig. 1E).

Recent evidences indicate that Smo signaling pathway inhibits neuronal apoptosis (Wang et al., 2018), whereas loss of Smo activity leads to increased neuronal apoptosis (Qin et al., 2019). In order to assess if Smo-CA or Smo-DN induces apoptosis in electroporated neurons, we performed immunohistochemical analysis for cleaved caspase-3, a marker for cell apoptosis (Logue and Martin, 2008) on electroporated rats at P15. Quantification of cleaved caspase-3-positive neurons shows no significant difference between the constitutively active mutant (0.24 ± 0.13 % and 0.34 ± 0.23 % in respectively Smo-CA and control; *p* = 0.70, Mann-Whitney test; Fig. 1F and G) or dominant negative (0.23 ± 0.15; *p* = 0.74, Mann-Whitney test; Fig. 1F and G) forms of Smo when compared to GFP control tissues.

Altogether, these observations indicate that expression of Smo-CA or Smo-DN does not alter the mean neuronal density and the positioning of pyramidal neurons in the cerebral cortex. Moreover, these results also suggest that IUE of Smo-related constructs does not affect apoptosis mechanisms in the postnatal cerebral cortex. By contrast, Smo-CA and Smo-DN change significantly the formation of synaptic terminals. These observations are in accordance with previous findings showing Smo-dependent enhancement of synapse formation in hippocampal neurons (Mitchell et al., 2012).

#### Smo-related constructs regulate the expression of its downstream target genes

To investigate the functional expression of the two Smo-related constructs, we performed immunofluorescence labeling by using antibody raised against Smo protein in somatosensory cortical tissues overexpressing Smo-related constructs at P15. As illustrated in figure 2A, the anti-Smo antibody detected well the expression of Smo-CA and Smo-DN in the electroporated somatosensory cortex. Smo activation mobilizes activator forms of Gli transcription factors, which in turn lead to an increase of Ptch1 and Gli1 expression (Jacob and Lum, 2007). To validate functional expression of Smo-related constructs, we quantified Gli1 and Ptch1 mRNA in cortices overexpressing Smo-CA or Smo-DN mutants using qRT-PCR. As expected, Gli1 mRNA expression are increased in Smo-CA when compared with GFP control animals (median values: 1.39 a.u. vs 0.76 a.u. in respectively Smo-CA and control; *p* = 0.041, Mann-Whitney test; Fig. 2B), and conversely decreased in Smo-DN (0.48 a.u.; *p* = 0.039 when compared to control, Mann-Whitney test; Fig. 2B). When compared with control or Smo-CA electroporated tissues, Ptch1 mRNA transcripts are downregulated in Smo-DN rats (3.61 a.u. in Smo-DN vs 6.97 a.u. in control and 6.21 a.u. in Smo-CA; *p* = 0.026 and *p* = 0.008 respectively, Mann-Whitney test; Fig. 2C). No significant difference is observed for Ptch1 mRNA between control and Smo-CA (6.21 a.u. for Smo-CA; *p* = 0.66, Mann-Whitney test; Fig. 2C). We next measured Shh protein expression levels in the somatosensory cortex of GFP, Smo-CA and Smo-DN at P15 and P30 by ELISA. Our results indicate that Shh protein is expressed at a constant and similar level in all samples analyzed (P15: 2.58 ng/mL for control vs 2.42 ng/mL for Smo-CA and 1.93 ng/mL for Smo-DN; *p* > 0.05; P30: 2.27 ng/mL for control vs 2.11 ng/mL for Smo-CA and 2.57 ng/mL for Smo-DN; *p* > 0.05, Mann-Whitney test; Fig. 2D).

**Figure 2:**
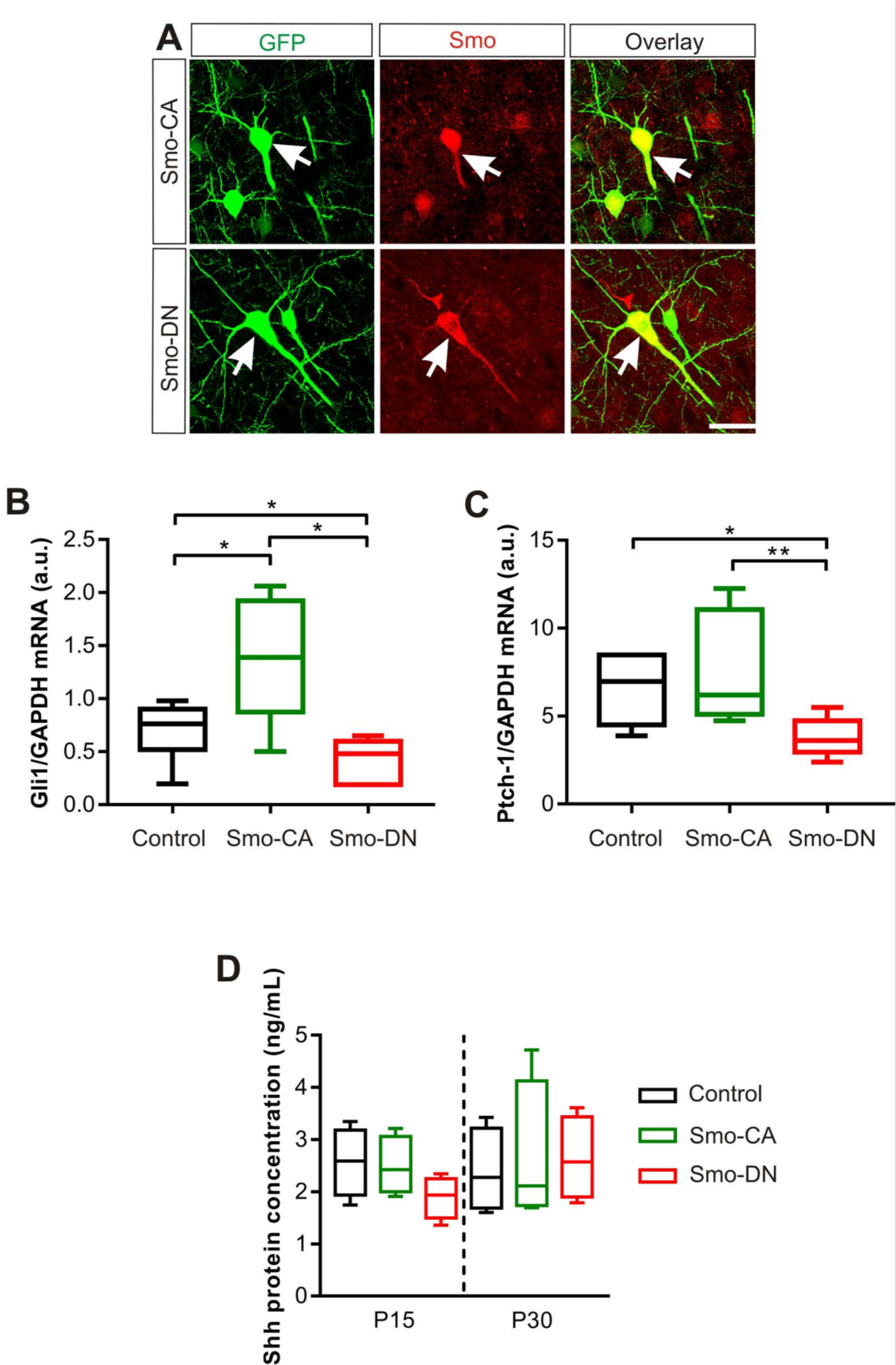
Smo-related constructs regulate the expression levels of Ptch1 and Gli1. (**A**) Cells overexpressing Smo related constructs (green) and immunostained with Smo antibody (red) showed a similar localization (overlay). Scale bar = 25 µm. (**B, C**) Box plots of relative expression of Gli1 and Ptch1 mRNA transcripts in GFP (control), Smo-CA and Smo-DN rats. (**B**) Box plots of Gli1 and (**C**) Ptch1 genes expression are normalized to GAPDH expression in the same RNA preparation. n = 6 rats for each condition. (**D**) Shh is expressed in post-natal somatosensory cortex. Box plots show median Shh protein concentration measured by ELISA at P15 and P30 in GFP (control), Smo-CA and Smo-DN from rat somatosensory cortex tissue lysate. n = 4 rats for each condition. \**p* > 0.05; ** *p* < 0.01.

These results confirm that in electroporated somatosensory cortices, the constitutively active form of Smo significantly promotes the expression of the target gene Gli1 whereas the dominant negative Smo acts in the opposite way by reducing its expression.

### Smo controls the developmental GABA excitatory to inhibitory shift

Cortical neuronal network construction during postnatal period requires the depolarizing to hyperpolarizing shift of GABA to occur at a precise timing during maturation (Ben-Ari and Holmes, 2005; Wang and Kriegstein, 2009; Wu and Sun, 2015). Because Shh signaling plays a nodal role in brain development (Álvarez-Buylla and Ihrie, 2014; Delmotte et al., 2020; Yao et al., 2016), we investigated whether the manipulation of Smo activity will affect the onset of GABA excitatory/inhibitory postnatal shift at the network level. We performed field recordings of Multi Unit Activity (MUA) in acute cortical slices from control (GFP), Smo-CA and Smo-DN rats and measured the effect of bath application of the GABA_A_R agonist isoguvacine (10 µM) on their spiking activity at P14, P20 and P30. Accordingly to the known depolarizing action of GABA in the immature neocortex (Kirmse et al., 2015; Riffault et al., 2018), isoguvacine induced an increase of spiking activity at P14 in cortical slices from control and Smo-DN rodents (median values: +33.87 %; *p* = 0.019 and +86.12 %; *p* = 0.019 respectively, Wilcoxon matched-pairs signed test; Fig. 3A and B). In contrast, isoguvacine decreased the spiking activity in Smo-CA mutants (−12 %; *p* = 0.009, Wilcoxon matched-pairs signed test, Fig. 3A and B). At P20 (Fig. 3B), isoguvacine induced either a decrease or an increase of spiking activity in control (−11.37 %; *p* = 0.519, Wilcoxon matched pairs signed test; Fig. 3B). In slices obtained from Smo-CA rats we observed a decrease of spikes frequency (−47.03 %; *p* = 0.011, Wilcoxon matched-pairs signed test) whereas in Smo-DN, isoguvacine still increased spikes frequency (+60.21%; *p* = 0.042, Wilcoxon matched-pairs signed test). At P30 (Fig. 3A-C), isoguvacine induced a decrease in spikes frequency from control and Smo-CA cortical slices (−22.86 % and −42.15 %; *p* = 0.008 and *p* = 0.015 respectively, Wilcoxon matched-pairs signed test), whereas it increases the spiking activity in Smo-DN rats (+12.89 %; *p* = 0.037, Wilcoxon matched-pairs signed test). Thus, the overall effects of isoguvacine on the spiking activity in the electroporated cortical slices suggest that the constitutively active and the dominant negative forms of Smo accelerate and delay respectively the developmental hyperpolarizing GABA shift (*p* < 0.0001, Chi-square test; Fig. 3C). In developing neocortex, the depolarizing-hyperpolarizing shift of GABA depends primarily on enhancement of the functional expression of KCC2 (Ben-Ari et al., 1989; Rivera et al., 1999). To investigate whether the hyperpolarizing shift of GABA observed in Smo-CA rats might involve activation of KCC2 we applied the KCC2-selective blocker VU0463271 (VU) on somatosensory cortex slices obtained from Smo-CA rats at P14. VU application shifted the isoguvacine response from inhibitory to excitatory when compared with non-treated Smo-CA slices (−12 % vs +57.1% with VU; *p* = 0.0009, Mann-Whitney test; Fig. 3D).

**Figure 3:**
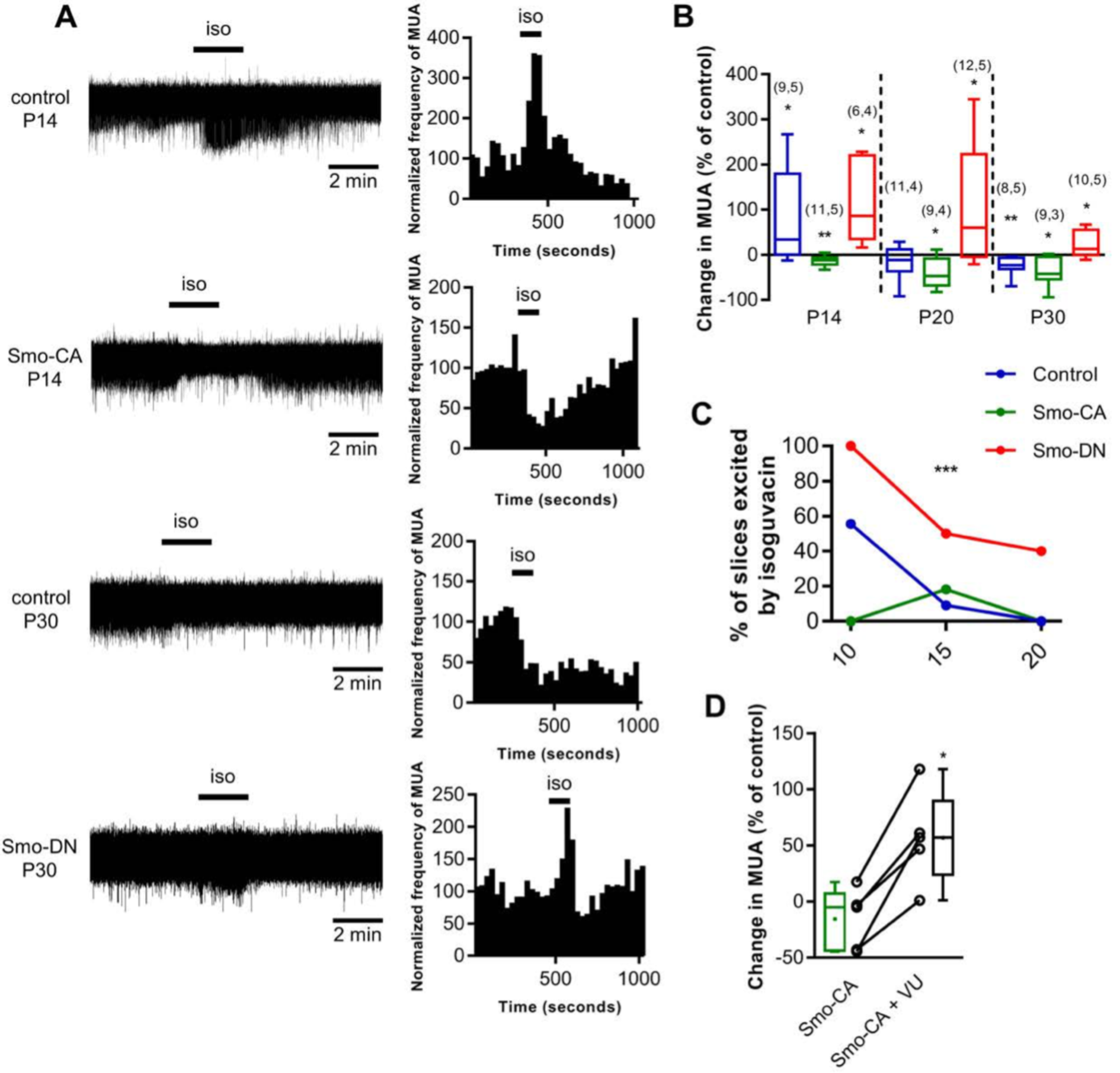
The developmental shift in the polarity of GABA_A_R mediated responses is modulated in cortical neurons expressing the dominant negative or the constitutively active forms of Smo. (**A**) Effects of isoguvacine (10 µM) in cortical slices from control GFP, Smo-CA and Smo-DN rats: representative traces of spontaneous extracellular field potentials (left column) recorded in electroporated area of cortical slices at P14 in GFP (control) and Smo-CA and at P30 in control and Smo-DN rats (left column). Corresponding time course of normalized frequency MUA is shown beside (right column) (**B**) Box plots of relative change of isoguvacine-dependent MUA frequency in electroporated rats at P14, P20 and P30. Numbers in parenthesis indicate the number of slices recorded and rats used. \**p* < 0.05; \*\**p* < 0.01 compared to control baseline; Wilcoxon test. (**C**) Developmental regulation of GABAergic excitation in electroporated rats. Proportion of excited slices is the proportion of slices showing an increase of 20% or more of MUA frequency during isoguvacine application. 3-5 rats per age and condition. \*\*\**p* < 0.001; Chi-square test. (**D**) Box plots of relative changes in MUA frequency during isoguvacine application with and without VU0463271 (VU) a KCC2-selective blocker application on Smo-CA rats at P14. n = 5 rats. \**p* < 0.05; Wilcoxon test.

Collectively, our data suggest that increasing Smo activity prematurely shifts GABA polarity from excitatory to inhibitory in a KCC2-dependent manner, whereas inhibiting Smo signaling delays the switch in GABA polarity.

#### Smo signaling controls chloride homeostasis

To further link Smo signaling with the shift in GABA polarity and neuronal chloride homeostasis, we performed gramicidin-perforated patch-clamp technique to measure the GABA_A_ reversal potential (E_GABA_) on dissociated hippocampal cultures in immature neurons (9 DIV). Consistent with our results obtained in cortical slices, the measurements of E_GABA_ in Smo-CA transfected neurons showed more hyperpolarized values when compared to control mCherry (−61.63 mV in Smo-CA vs −50.13 mV in control *p* = 0.008, Mann-Whitney test; Fig. 4A and B). Contrary to action of Smo-CA, the overexpression of Smo-DN did not produce statistically significant change of E_GABA_ as compared to control mCherry transfected neurons (–57.13 mV in Smo-DN; *p* = 0.38, Mann-Whitney test; Fig 4B).

**Figure 4:**
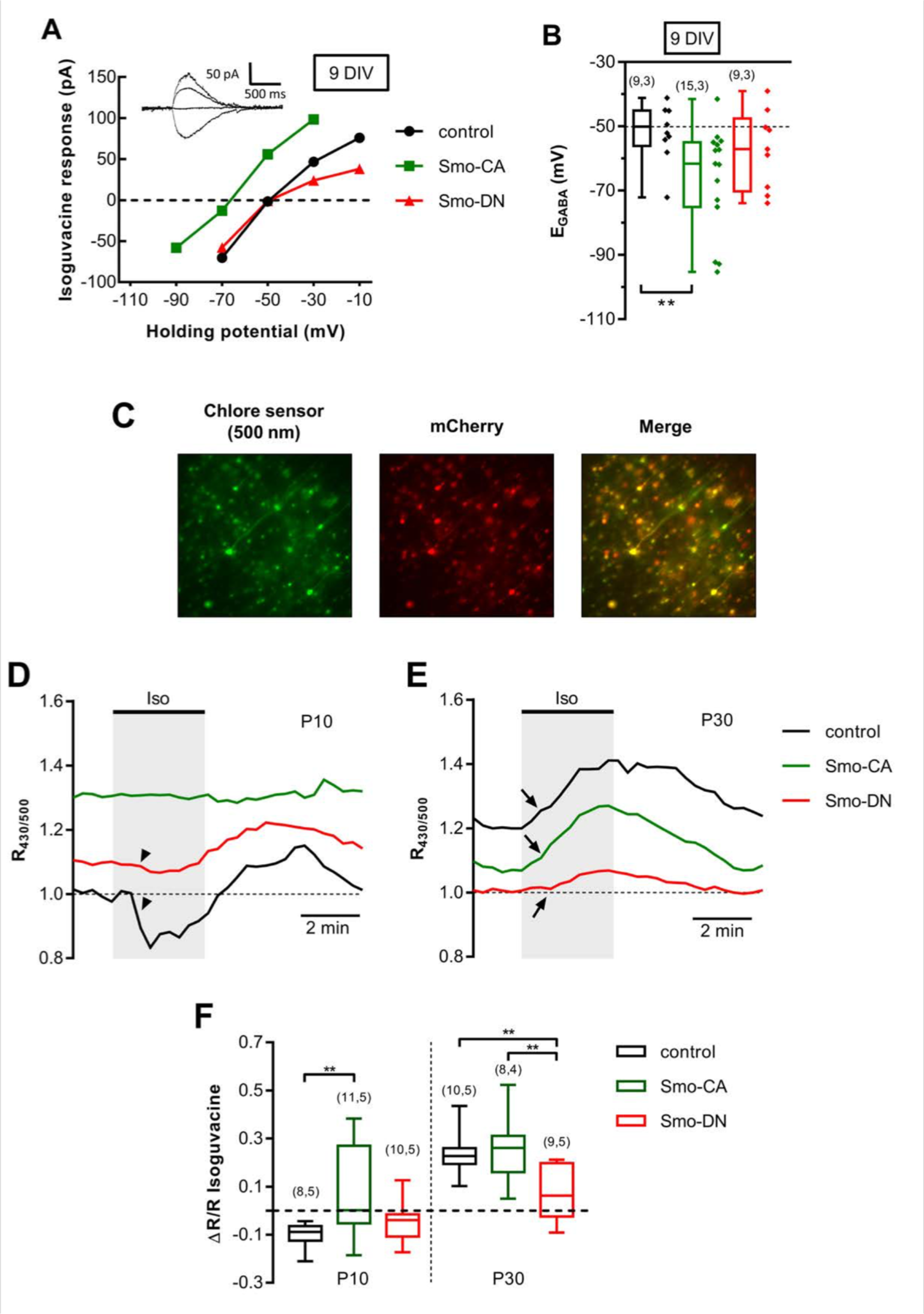
Smo signaling controls chloride homeostasis in cultured neurons and acute slices. (**A**) Gramicidin perforated patch clamp recordings of I-V relationships for isoguvacine currents in mCherry only (control) or mCherry plus Smo-CA or Smo-DN in rat hippocampal primary neuronal cultures (9 DIV). Inserts depict the isoguvacine currents for control condition. Scale bar, 500 ms, 50 pA. (**B**) Box plots of E_GABA_ in the indicated conditions. The number of cells recorded and cultures used are indicated in parenthesis. (**C**) Typical images of Cl-Sensor fluorescence excited at 500nm in slice from P30 rat electroporated with Cl-Sensor plus Smo-CA. Rats were electroporated *in utero* at embryonic day 15 with Cl-Sensor plus mCherry (control) or Cl-Sensor plus Smo-CA (Smo-CA) or Cl-Sensor plus Smo-DN (Smo-DN). The regions of interest were drawn around the soma of cells located in the focal plane. 5 to 10 neurons were analyzed per image. Data represent results obtained from 4-5 rats per experimental condition and 1-2 slices were recorded per animal. (**D and E**) Example of typical ratiometric fluorescence (R430/500) recordings at P10 (**D**) and P30 (**E**) from control, Smo-CA and Smo-DN rats. Horizontal bars indicate the times of application of ACSF containing isoguvacine. Arrows and arrowheads indicate different types of responses between control, Smo-CA and Smo-DN neurons at P10 and P30. Data represent results obtained from 4-5 rats per experimental condition and 1-2 slices were recorded per animal. (**D**) Box plots quantification of fluorescence change of ΔR/R, where R is the mean of 5 measurements before isoguvacine application and ΔR is the difference between absolute maximum of isoguvacine-induced response. Number of slices and rats are indicated in parenthesis. \**p* < 0.01, \*\**p* < 0.01 when compared to control at the same age; Mann-Whitney U test.

To corroborate the results obtained in neuronal cultures we studied whether Smo signaling modifies [Cl^-^]_i_ in acute slices of somatosensory cortex isolated from Smo-CA and Smo-DN rats. For this purpose, mCherry control, Smo-CA or Smo-DN were co-electroporated *in utero* at E15 with constructs encoding ratiometric Cl-sensor (Friedel et al., 2013; Friedel et al., 2015) (Fig. 4C). Electroporated slices prepared at postnatal days P10 and P30 harbored hundreds of neurons expressing Cl^-^-Sensor in cortical layers V/VI. The bath application of isoguvacine (10 µM, 3 min) to P10 Smo-DN and control slices produces a uniform decrease of R_430/500_ (indicated by an arrowhead in Fig. 4D and by the negative value of ΔR/R in Fig. 4F), reflecting a chloride ion extrusion characteristic of depolarizing action of GABA (median values: −0.04 a.u. for Smo-DN vs −0.09 a.u. for control; *p* = 0.17, Mann Whitney test; Fig. 4B and D), whereas in Smo-CA slices, isoguvacine triggered either an increase, no change or slight decrease of R_430/500_. The median of these changes in Smo-CA slices was +0.00 a.u. and statistical analysis revealed significant difference when compared to control (*p* = 0.005, Mann Whitney test; Fig. 4D and F). This difference indicates that in Smo-CA slices GABA is switching from depolarizing to hyperpolarizing actions. In P30 control and Smo-CA slices, isoguvacine produced a chloride influx (indicated by an arrow in Fig. 4E) as shown by the positive value of ΔR/R (+0.22 a.u. for control vs +0.26 a.u. for Smo-CA; *p* = 0.63, Mann-Whitney test; Fig. 4E and F), whereas Smo-DN slices showed a slight increase of R_430/500_ (+0.06 a.u.; *p* = 0.002 and *p* = 0.01 when compared to control and Smo-CA respectively; Fig. 4E and F). The latter finding suggest that Smo-dependent pathway is active at rest in mature P30 neurons to maintain the low [Cl^-^]_I_ and hyperpolarizing inhibitory action of GABA. The inhibition of Smo action using Smo-DN leads to increase [Cl^-^]_I_ that reduces inhibitory strength of GABA. Thus, in both immature acute slices and primary cultures, activation of Shh using Smo-CA lead to hyperpolarizing shift of Cl^-^, whereas Smo-DN did not affect the [Cl^-^]_I_ at this stage.

### Smo regulates the activity of KCC2

The formation of KCC2-dependent developmental shift of GABA rely on complex mechanism involving progressive increase of the amount of KCC2 protein and its posttranslational modifications via at least two distinct phosphorylation sites, serine 940 (Ser^940^) and threonine 1007 (Thr^1007^) (Côme et al., 2019; Medina et al., 2014; Moore et al., 2017). We measured the protein and mRNA expression of KCC2 from the somatosensory cortices of Smo-CA and Smo-DN rats at P10. We have observed similar levels between control and Smo-related constructs for KCC2 mRNA (median values: 5.02 a.u. for control tissues vs 3.3 a.u. for Smo-CA, *p* = 0.84; and 2.92 a.u. for Smo-DN, *p* = 0.42 compared to control; Mann-Whitney test; Fig. 5A) and proteins (1.23 a.u. for control tissues when normalized to ß3 tubulin vs 1.26 a.u. for Smo-CA and 1.25 a.u. for Smo-DN; *p* = 0.48 and *p* = 0.48 respectively, Mann Whitney test; Fig. 4B and C). The membrane stability and transporter activity of KCC2 are dependent on the phosphorylation state of intracellular C-terminus domains. For instance, phosphorylation of the Ser^940^ residue increases stability and functionality of KCC2, whereas phosphorylation on the Thr^1007^ residue enhances KCC2 endocytosis (Lee et al., 2007; Medina et al., 2014). To determine which steps in KCC2 trafficking may be regulated by Smo-related constructs, we carried out quantitative immunoblotting assay using antibodies recognizing phosphorylated forms of Ser^940^ (Lee et al., 2011) and Thr^1007^ (de Los Heros et al., 2014). We found that the ratio of phospho-Ser^940^ (pKCC2 S^940^) to total KCC2 protein (pKCC2 S^940^/KCC2) is upregulated in Smo-CA electroporated cortices when compared to control and Smo-DN conditions (0.56 a.u. for Smo-CA vs 0.39 a.u. for control and 0.27 a.u. for Smo-DN; *p* = 0.02 and *p* = 0.02 respectively compared to Smo-CA, Mann-Whitney test; Fig. 5B and D). The ratio of pKCC2 Thr^1007^ to total KCC2 protein (pKCC2T1007/KCC2) was not modified in Smo-related constructs (0.90 a.u. for control tissues vs 0.89 a.u. for Smo-CA and 0.78 a.u. for Smo-DN; *p* = 0.88 and *p* = 0.17, when compared to control respectively, Mann-Whitney test; Fig. 5B and E).

**Figure 5:**
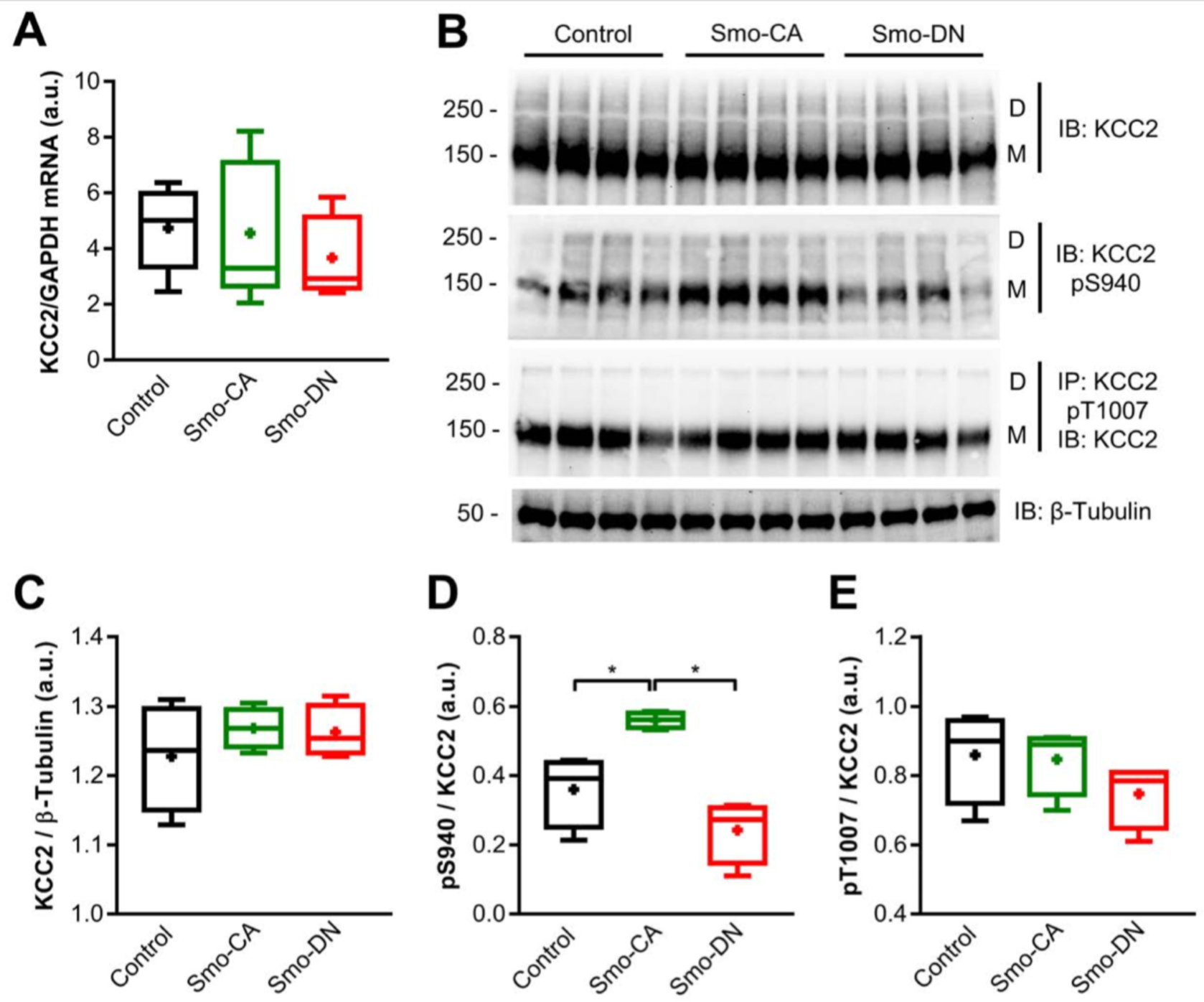
KCC2-Ser^940^ phosphorylation is regulated by Smo signaling. (**A**) Box plots of relative expression of KCC2 mRNA normalized to GAPDH mRNA level in P10 rats electroporated with GFP alone (Control), GFP + Smo-CA (Smo-CA) or GFP + Smo-DN (Smo-DN). N = 5 rats for each condition. (**B**) Immunoblots for total KCC2 and phosphorylated forms of KCC2 Ser^940^ (KCC2 pS940) and KCC2 Thy^1007^ (KCC2 pT1007) in protein extracts from P10 electroporated somatosensory cortices. Molecular weights were indicated on the left of the blot (in kDa). (**C, D and E**) Box plots of normalized total KCC2(**C**); KCC2 pSer^940^ (**D**), KCC2 pThr^1007^ (**E**) relative protein level. n = 4 rats for each condition. *p < 0.05; Mann-Whitney U test.

Together, these data suggest that enhanced Smo activity led to KCC2 post-translational modifications, with increased Ser^940^ phosphorylation in Smo-CA electroporated somatosensory cortex.

#### The dominant-negative form of Smo affects the stability of KCC2pH_ext_ to the plasmalemmal surface by promoting endocytosis

The phosphorylation of Ser^940^ described above is well known for its ability to stabilize KCC2 in the plasma membrane whereas dephosphorylation of Ser^940^ promotes KCC2 endocytosis (Kahle et al., 2013; Lee et al., 2007). We therefore assessed whether Smo-related constructs are able to regulate KCC2 trafficking to the cell surface. As tool we used a KCC2 construct tagged in an external loop with a fluorescent protein pHluorin (KCC2-pH_ext_) (Kahle et al., 2014). This construct allows both measurement of KCC2-pH_ext_ -dependent shift of [Cl^-^]_i_ and visualization of surface expression/internalization of KCC2-pH_ext._ (Friedel et al., 2015; Friedel et al., 2017).

As expected, the overexpression of KCC2-pH_ext_ in 9 DIV neurons resulted in a negative shift of E_GABA_ (median values from −50.13 mV to −86.25 mV; *p* < 0.0001, Mann-Whitney test; Fig. 6A and B), indicating also that at least some portion of KCC2-pH_ext_ molecules were delivered to the plasma membrane. The overexpression of Smo-CA did not produce additional hyperpolarizing shift of E_GABA_, in KCC2-pH_ext_ -positive neurons (−83.82 mV for Smo-CA + KCC2-pH_ext_ vs −86.25 mV for mCherry + KCC2-pH_ext_; *p* = 0.69, Mann-Whitney test; Fig. 6A and B) whereas overexpression of Smo-DN resulted in a significant depolarizing shift of this parameter (−53.1 mV in Smo-DN + KCC2-pH_ext_ vs −86.25 mV in mCherry+ KCC2-pH_ext_; *p* = 0.006, Mann-Whitney test; Fig. 6A and B) indicating on either Smo-DN-dependent inactivation of the membrane pools of KCC2-pH_ext_ or reduction of its expression in plasma membrane.

**Figure 6:**
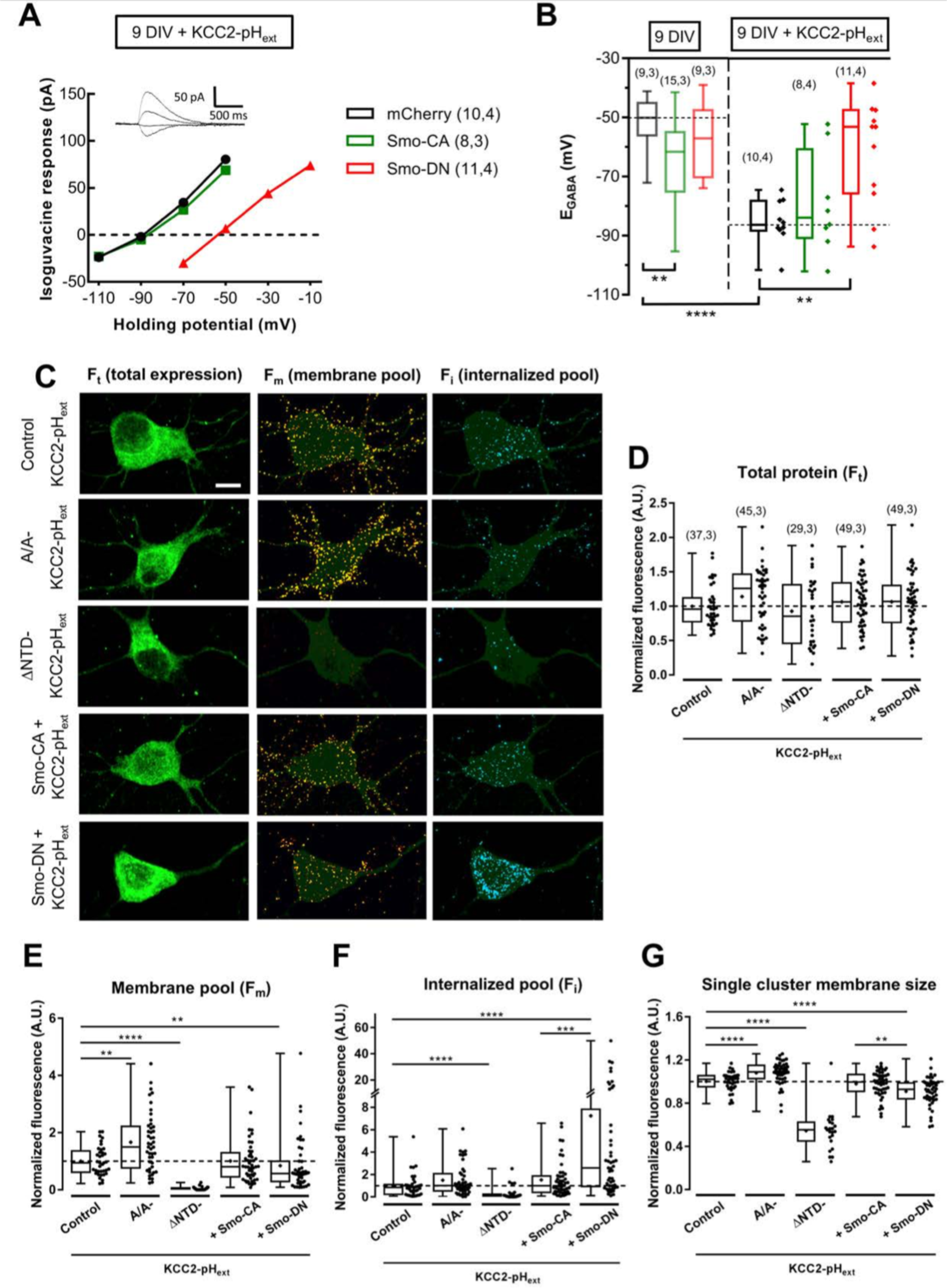
Dominant negative form of Smo affects the stability of KCC2 to the plasmalemmal surface. (**A**) Gramicidin perforated patch clamp recordings I-V relationships for isoguvacine currents in KCC2-pH_ext_ alone or co-transfected with Smo-CA (green) or Smo-DN (red) in hippocampal primary culture at 9 DIV. Inserts depict the isoguvacine currents for control condition. Scale bar, 500 ms, 50 pA. (**B**) Box plots of E_GABA_ in the indicated conditions. Left panel show neurons transfected without KCC2-pH_ext_ from data shown in Fig. 4A and B. Right panel show neurons co-transfected with KCC2-pH_ext_. The number of cells recorded, and cultures used are indicated in parenthesis. (**C**) Representative images illustrating total, membrane and internalized pools of KCC2 with external tag (KCC2-pHext) in vehicle (control), and Smo-related transfected constructs in hippocampal primary culture neurons. KCC2 construction stable to membrane (A/A KCC2) was used as positive control for KCC2 membrane trafficking. Neurons expressing a KCC2 mutant construction known to not be addressed to membrane (ΔNTD-KCC2) were proceeded in parallel experiments to ensure that immunocytochemistry on living neurons does not permeabilized the membrane. Scale bar 20μm. (**D**) Box plots of total protein (F_t_), (**E**) normalized membrane (F_m_), (**F**) internalized pool (F_i_) fluorescence and (**G**) single membrane cluster size in cultured neurons expressing the indicated constructs. The number of cells and cultures used indicated in parenthesis are identical for all box plots. The sample size is the same for all box plots. \**p* <0.05, \*\**p* < 0.01, \*\*\**p*<0.005, \*\*\*\**p*<0.001; Mann-Whitney U test.

To further corroborate this study, we performed a live-staining analysis of surface expressed/internalized KCC2-pH_ext_. Three different pools of KCC2-pH_ext_ were revealed using multistep immunolabeling protocol: the total expressed amount of protein (F_t_), the amount of KCC2-pH_ext_ cell surface expression (F_m_) and the amount of KCC2-pH_ext_ internalized (F_i_) (Fig. 6C). As positive and negative controls we used overexpression of KCC2-pH_ext_ mutants T906A/T1007A (A/A-KCC2-pH_ext_) and ΔNTD-KCC2-pH_ext_ known for their increased and perturbed surface expression abilities, respectively (Friedel et al., 2017). In the performed experiments, the live-cell immunolabelling of the control non-mutated KCC2-pH_ext_ revealed well detectable F_m_ signal in form of clusters, whereas the post-hoc multistep immunolabelings revealed F_i_ pool of molecules and total amount of expressed KCC2-pH_ext_ (Fig.6C). The relative amounts of F_m_ and membrane labeled cluster size in KCC2-pH_ext_ neurons were lower than those in A/A-KCC2-pH_ext_ expressing neurons and significantly higher that the values revealed in ΔNTD-KCC2-pH_ext_ – transfected cells (median values: For F_m_: 0.94 a.u. for KCC2-pH_ext_ vs 1.49 a.u. for A/A-KCC2-pH_ext_ and 0.008 a.u. for ΔNTD-KCC2-pH_ext_; *p* = 0.002 and *p* < 0.0001, respectively, Fig. 6E; For single cluster membrane size: 1.02 a.u. for KCC2-pH_ext_ vs 1.09 a.u. for A/A-KCC2-pH_ext_ and 0.55 a.u for ΔNTD-KCC2-pH_ext_; *p* < 0.0001 and *p* < 0.0001, respectively; Mann-Whitney test; Fig. 6G). The co-expression with KCC2-pH_ext_ of Smo-CA did not produce statistically significant change in F_i_, F_m_ or single surface labeled cluster size (Fig. 6E, F and G). By contrast, neurons co-expressing KCC2-pH_ext_ and Smo-DN showed a significant twofold higher internalization rate of labeled KCC2-pH_ext_ molecules (0.84 a.u. for KCC2-pH_ext_ vs 2.51 a.u. for Smo-DN + KCC2-pH_ext_; *p* < 0.0001; Fig. 6C and F), significant twofold lower amount of F_m_ (0.94 a.u. for KCC2-pH_ext_ vs 0.57 a.u. for Smo-DN + KCC2-pH_ext_; *p* = 0.009; Mann-Whitney test; Fig. 6C and E) and significantly smaller size of surface located KCC2-pH_ext_ clusters (1.02 a.u. for KCC2-pH_ext_ vs 0.92 a.u. for Smo-DN + KCC2-pH_ext_; *p* < 0.0001, Mann-Whitney test; Fig. 6G).

Thus, consistent with data illustrated in Figures 3-6, both [Cl^-^]_i_ measurement and live-cell immunolabelling revealed a striking Smo-dependent change in functioning and surface expression of KCC2-pH_ext_.

### Smo-DN rats display increased susceptibility to PTZ-induced seizures

Large number of studies have illustrated that the KCC2 dysfunction facilitates initiation of epileptic seizures (Chen et al., 2017; Moore et al., 2018). The changes of KCC2’s Ser^940^ phosphorylation that modify neuronal chloride homeostasis and depolarizing strength of GABA strongly affect the seizures susceptibilities in KCC2 transgenic mice (Silayeva et al., 2015). We finally investigated whether overexpression of Smo mutants may influence the susceptibility to seizures. Smo-DN, Smo-CA and age-matched control rats (transfected with GFP alone) at P30 were intraperitoneally injected with subconvulsive doses of PTZ (25 mg/kg; Fig. 7A), an inhibitor of the GABA_A_ receptors (Klioueva et al., 2001). Previous studies showed that the lowest threshold susceptibility to PTZ can be observed during the first 2 postnatal weeks and the highest at around P30 (Klioueva et al., 2001; Vernadakis and Woodbury, 1969). These results suggest that the vulnerability to PTZ is correlated to the maturation of GABAergic system during postnatal development. Injections of PTZ were continued every 10 min until each rat had generated generalized tonic-clonic seizure. Seizure susceptibility and severity were quantified with the canonical Racine’s scale of seizure severity (Lüttjohann et al., 2009), including dose and latency to onset of the first generalized tonic-clonic seizure. Generalized seizures in Smo-DN rats occur at lower doses of PTZ (median values: 87.5 mg/kg compared with 100 mg/kg for both control and Smo-CA rats; *p* = 0.005 and *p* = 0.003 respectively, Mann-Whitney test; Fig. 7C) and with shorter latencies than control and Smo-CA rats (30.83 min compared with 32.71 min in control and 32.69 min in Smo-CA; *p* = 0.041 and *p* = 0.024 respectively; Mann-Whitney test; Fig. 7B). No significant difference is observed between control and Smo-CA rats in the latency (32.69 min compared with 32.71 min in control; *p* = 0.5158; Fig. 7B) or cumulative dose of PTZ to induce tonic-clonic seizures (*p* = 0.501; Fig. 7C).

**Figure 7:**
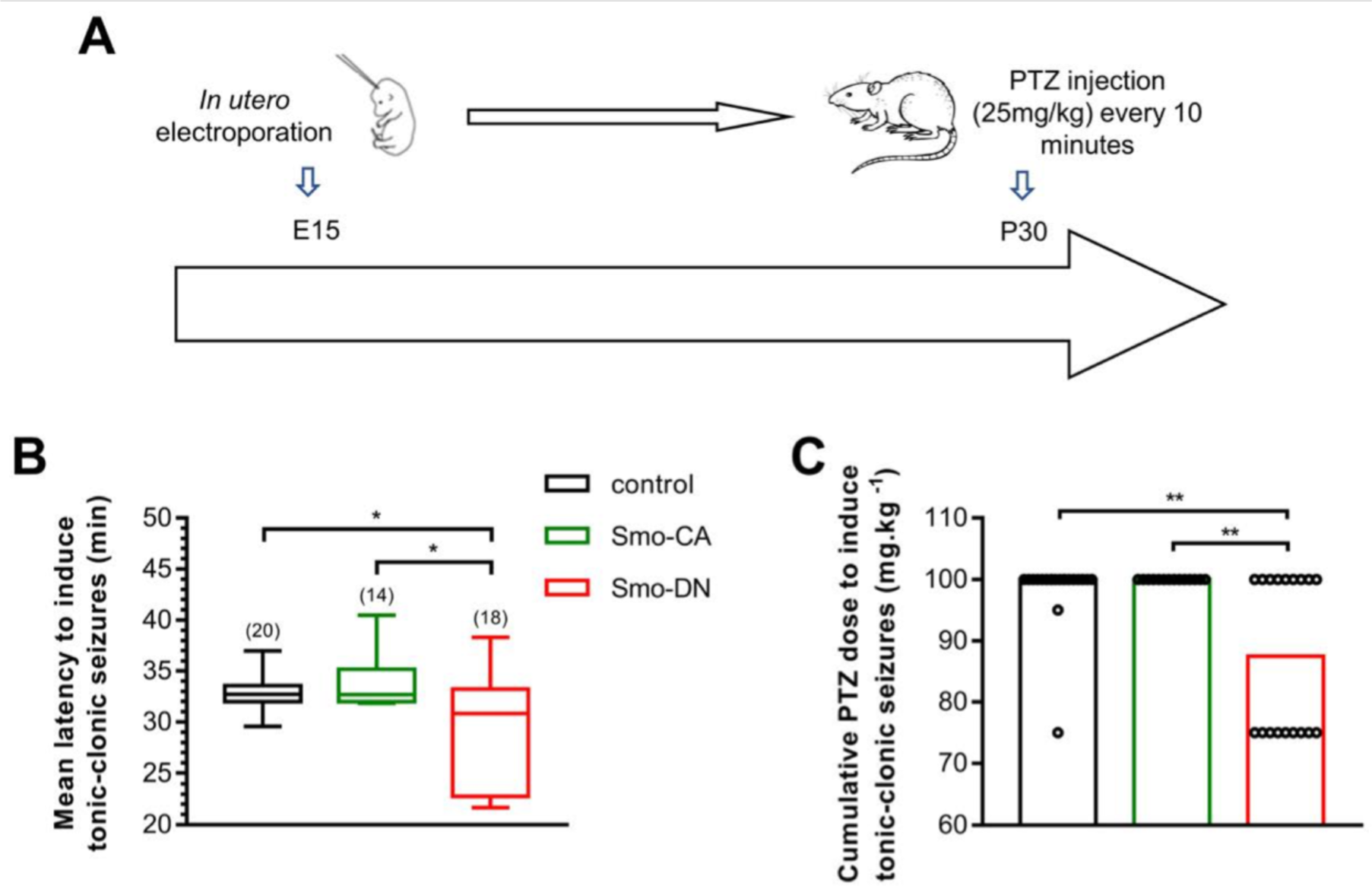
Modified susceptibility to pentylenetrazol-induced (PTZ) seizures in Smo mutant rats. (**A**) Experimental paradigm design. (**B**) Box plots of latency needed to induce generalized tonic-clonic seizures in 3 groups of electroporated rats: control (electroporated with GFP), Smo-CA and Smo-DN. (**C**) Bar graph of PTZ cumulative dose needed to induce generalized tonic-clonic seizures. Bars are medians with experimental points. Numbers in parenthesis indicate the number of rats used. \**p* < 0.05; \*\**p* < 0.01; Mann-Whitney U test.

These results indicate that reduced-seizure threshold in Smo-DN rats could be due to a downregulation of KCC2 and deregulation of chloride homeostasis induced by the diminution of Smo activity.

## Discussion

In the present study, we investigated the actions of Shh-Smo signaling pathway on GABAergic transmission *in vivo* and *ex vivo* in the rat somatosensory cortex and *in vitro* in primary culture of hippocampal neurons using dominant-negative (Smo-DN) and constitutively active (Smo-CA) forms of Smo. We show that activation of Smo signaling in immature neurons at early postnatal stages accelerated the process of GABAergic switch form depolarizing to hyperpolarizing activity in a KCC2-dependent manner, leading to a low intracellular chloride concentration. Conversely, Smo signaling blockade altered the phosphorylated state of KCC2 and membrane stability, weakening the GABAergic inhibition on later stages of the development, and resulting in an increase of seizure susceptibility. These findings reveal an unexpected function of the G-protein-coupled receptor Smoothened on chloride homeostasis regulation and, in particular, on the timing of the GABA shift from depolarizing-to hyperpolarizing in the postnatal developing brain.

The postnatal developmental switch in GABA polarity is tightly regulated by the actions of several peptides including neurotrophic factors (Aguado et al., 2003; Kelsch et al., 2001; Ludwig et al., 2011; Riffault et al., 2018), hypothalamic neurohormones (Leonzino et al., 2016; Spoljaric et al., 2017; Tyzio et al., 2006), central and peripheral hormones (Dumon et al., 2018; Sawano et al., 2013). Hence, the precise timing of GABAergic polarity shift depends on the balance between factors promoting or delaying the onset of GABAergic inhibitory transmission. It is noteworthy that these factors are developmentally regulated and/or operate within specific time windows to control the functional expression of KCC2 and associated chloride homeostasis and, consequently the postnatal maturation of GABAergic transmission in the central nervous system. Interestingly, the excitatory action of GABA at early stages of the development coincide with the predominance of trophic factors responsible for the inhibition of KCC2 expression and function, such the immature form of BDNF (proBDNF) (Riffault et al., 2018) and the adipocyte hormone leptin (Dumon et al., 2018). These two factors are expressed shortly after birth and surge during the first postnatal week in rodents (Dumon et al., 2018; Menshanov et al., 2015). They have been shown to promote a low KCC2 protein and mRNA expression levels and consequently a high intracellular chloride concentration, thus maintaining a depolarizing action of GABA in neonates. The developmental decreases of leptin and proBDNF observed at the end of the first postnatal week in rodents, are concomitant to an increase in oxytocin (Leonzino et al., 2016; Tyzio et al., 2006), mature BDNF (mBDNF) (Maisonpierre et al., 1990) and thyroid hormones (Sawano et al., 2013), which in turn, up-regulate KCC2, leading to a low intracellular chloride and a hyperpolarizing action of GABA. Although we did not observe any difference in the level of Shh proteins between P15 and P30, a study demonstrated that Shh expression paralleled the temporal expression of mBDNF in the cortex of rodents with a nearly undetectable level at birth and a continuous increased expression at both mRNA and protein levels from during the two first postnatal weeks (Rivell et al., 2019). This temporal expression of Shh and its signaling partners during postnatal brain development has been shown to play a role in neuronal network construction through axonal growth and synaptic maturation (Yao et al., 2015). Here, we uncover a new critical function of Shh components in the regulation of the GABA polarity switch. Indeed, we showed that activation of the GABA_A_ receptor by isoguvacine, in immature (i.e. post-natal day 14) Smo-CA cortical neurons results in an influx of chloride and a decrease in spiking activity whereas, in mature (i.e. post-natal day 30) dominant negative Smo cortical neurons (Smo-DN), maintains a depolarizing and excitatory action of GABA. Thus, in the absence of Smo signaling, the GABA shift is delayed, thereby leading to an altered chloride homeostasis and higher susceptibility to seizures, presumably reflecting an unbalanced excitation to inhibition (E/I) ratio. E/I imbalance has been linked to cognitive disorders such as schizophrenia, autism spectrum disorders (ASD), depression and epilepsy (Ben-Ari and Holmes, 2005; Brambilla et al., 2003; Kuzirian and Paradis, 2011; Mueller et al., 2015). Abnormal levels of Shh have been also observed in ASD patients with either an increased serum level in children or a decreased mRNA level in post-mortem adult human brain tissue from autism patients (Choi et al., 2014; Halepoto et al., 2015). In addition, mutations in Ptchd1, a gene displaying secondary structures similar to Ptch1, are found in patients with intellectual disability and ASD (Ung et al., 2018). Consistent with a possible role of Shh components in the regulation of the E/I balance, these observations also suggest that an alteration in Shh-Ptch-Smo pathway may have an implication to the pathophysiological mechanisms in ASD. In line with these observations, the role of Shh signaling in neuronal activity has been recently addressed in an elegant study (Hill et al., 2019) showing that the selective disruption of Shh signaling with a conditional knockout of Smo leads to an increase in neuronal excitability of cortical neurons, thus highlighting the importance of Shh on the E/I equilibrium and the construction of functional cortical networks. The results from this study are consistent with our findings showing that Shh-Smo signaling blockade increases neuronal network excitability in rodents expressing the dominant negative form of Smo. Furthermore, we found that blockade of Smo activity in Smo-DN rodents leads to a decrease in KCC2 function and to a reduced threshold for PTZ-induced tonic-clonic seizure. In agreement with a previous study by Chen et al. (Chen et al., 2017), these results suggest that downregulation of KCC2 function may contribute to epileptic seizures. The implication of Shh signaling in epilepsy and seizures has been indirectly addressed in previous studies showing that somatic mutations in Shh pathway genes in human are associated with hypothalamic hamartoma and drug-resistant epilepsy (Hildebrand et al., 2016; Saitsu et al., 2016). Likewise, the Shh expression was increased in hippocampus and neocortex from both human and animal models of temporal lobe epilepsy (TLE) (Fang et al., 2011), thus suggesting a potential relationship between epilepsy and Shh activity. In line of the above studies, Feng et al. (Feng et al., 2016) reported that during TLE development, Shh contributes to epileptogenesis through the inhibition of glutamate transporters function leading to abnormal extracellular glutamate levels, whereas the blockade of Shh signaling pathway reduced the severity of seizures. Together, these observations can lead to opposing interpretations regarding the possible interplay between Shh pathway and epilepsy. These apparent discrepancies could be explained by the treatments utilized (i.e. acute seizure models with PTZ versus spontaneous seizures in the pilocarpine model of chronic TLE) that are able to activate different downstream signaling cascades (Choudhry et al., 2014). Moreover, other studies have found that the Shh-Smo/Gli pathway activation increased the synthesis and secretion of neurotrophic factors like nerve growth factor (NGF) and BDNF (Bond et al., 2013; Chen et al., 2018; Radzikinas et al., 2011), which they may have neuroprotective effects in TLE (Bovolenta et al., 2010; Falcicchia et al., 2018; Paradiso et al., 2009). Clearly, additional studies are needed to get a deeper knowledge on the cellular and molecular mechanisms linking epilepsy to Shh pathway components.

Importantly, our *in vitro* experiments revealed that in immature neurons of both primary rat hippocampal cultures and rat neocortical acute slices the E_GABA_ shifted towards a more hyperpolarized level in Smo-CA expressing neurons and indicating that Smo-CA initiated decrease of neuronal [Cl^-^]_i_. In immature neurons the increased [Cl^-^]_i_ and correspondent depolarizing values of E_GABA_ are determined primarily by effective intrusion of Cl^-^ by sodium/potassium/chloride cotransporter type 1 (NKCC1) and absence of the effective Cl^-^ extrusion by KCC2. The blockage of NKCC1 (Yamada et al., 2004) or activation of KCC2 (Dumon et al., 2018; Inoue et al., 2012; Khirug et al., 2005) lead both to hyperpolarizing shift of E_GABA_. The present study revealed a novel Smo-dependent pathway of KCC2-upregulation during neuronal development. Whether Smo pathway regulates also NKCC1 activity as well as other transporters and channels controlling Cl-homeostasis in immature neurons should be a subject of independent respective projects.

The Cl-extrusion activity of KCC2 in developing neurons depends on the amount of protein expression (Rivera et al., 1999) and its post-translational modifications that regulate the KCC2’s surface targeting and clustering (Côme et al., 2019; Medina et al., 2014). Thus, KCC2 cycle between synaptic sites and endocytic compartments which depends on the phosphorylation and/or dephosphorylation state of different amino-acid residues (Côme et al., 2019; Kahle and Delpire, 2016; Kahle et al., 2014), but also this membrane turnover requires regulation to its cytosolic N-and C-termini (Friedel et al., 2017). For instance, N-terminal truncation of KCC2 prevents its export at the cell surface, whereas C-terminal truncation leads to a decreased surface expression of KCC2 (Friedel et al., 2017). The observed loss of KCC2 surface expression in the C-terminal mutants is explained by an increase in the internalization rate without affecting cell surface delivery. Furthermore, KCC2 function can also be regulated through lateral diffusion at the plasma membrane (Chamma et al., 2013). This suggests that KCC2 membrane dispersion may, in turn, mediate membrane destabilization and endocytosis resulting in intracellular chloride homeostatic dysregulation (Côme et al., 2019). In agreement with this, we found that inhibition of Smo signaling pathway in mature neurons induces a decrease in KCC2 cluster size and enhances the intracellular concentration of chloride that sets E_GABA_ at more positive values than the resting potential with a reduced plasmalemmal levels of KCC2 cell surface expression levels. We further show that these effects occur through an increased rate of KCC2 turnover.

In conclusion, these data uncovered an unexpected role of Shh-Smo components during early postnatal development through the control of chloride homeostasis. Thus, Smo signaling will tune up or down GABA inhibitory transmission by controlling phosphorylation of KCC2 that affects its stability in the cell membrane and modulates neuronal chloride homeostasis. Finally, these results suggest that Smo signaling is able to set the cursor of GABAergic inhibitory switch during this critical maturational period where developmental pathogenesis takes place.

## Material and methods

All animal procedures were carried out according to the guidelines set by the INSERM animal welfare through the local committee (CEEA n°14) and the European Communities Council Directives (2010/63/UE). Male and female Wistar rats were purchased from Janvier Labs (https://www.janvier-labs.com/fiche_produit/rat_wistar/). Animals were raised and mated at INMED A2 animal facility housed under a 12 h light/dark cycle at 22-24°C and had access to food and water *ad libitum*.

### Reagents and treatments

The following reagents were purchased from the indicated sources: 1,2,3,4-Tetrahydro-6-nitro-2,3-dioxo-benzo[f]quinoxaline-7-sulfonamide (NBQX) was obtained from the Molecular, Cellular and Genomic Neuroscience Research Branch (MCGNRB) of the National Institute of Mental Health (NIMH, Bethesda, MD, USA). TTX was purchased from abcam (Cambridge, UK). Isoguvacine, bicuculline and VU0463271 were purchased from Tocris Cookson (Bristol, UK). Strychnine and Bumetanide from Sigma (St-Louis, Missouri, USA).

### In utero electroporation

*In utero* injections and electroporations were performed in embryos from timed pregnant rats (embryonic day 15) that received buprenorphine (Buprecare at 0.03 mg/kg) and were anaesthetized with sevoflurane (4.5%) 30 minutes later (Martineau et al., 2018). Briefly, the uterine horns were exposed, and a lateral ventricle of each embryo was injected using pulled glass capillaries and a microinjector (PV 820 Pneumatic PicoPump; World Precision Instruments, Sarasota, FL) with Fast Green (2 mg/ml; Sigma, St Louis, MO, USA) combined with the following DNA constructs encoding GFP or mCherry and/or Smo-DN and/or Smo-CA and/or Cl-Sensor (ratio 1:2). Plasmids were further electroporated by delivering 40V voltage pulses with a BTX ECM 830 electroporator (BTX Harvard Apparatus, Holliston, MA, USA). The voltage was discharged in five electrical pulses at 950 ms intervals via tweezer-type electrodes (Nepa Gene Co, Chiba, Japan) placed on the head of the embryo across the uterine wall. We performed *in utero* electroporation in embryonic rats at E15, corresponding to an active period of both radial and tangential migration of newborn neurons in the cortex (Kriegstein and Noctor, 2004). At birth, successfully electroporated pups were selected after transcranial visualization of the GFP reporter fluorescent proteins only. Following experimentation, morphological analysis of electroporated tissues, selected by GFP or mCherry expression under fluorescent stereomicroscope (Olympus SZX 16), was performed. The criteria commonly used for selection of animals were localization and cell density of transfected cortices and absence of abnormal cortex morphology (i.e. cortical disruption, thickening and abnormal cortical organization). Our analyses revealed no alterations in cortical layer position or morphology in GFP or mCherry, Smo-CA and Smo-DN conditions, although we cannot exclude the possibility that Smo constructs impact the neuronal cells at the cellular and subcellular levels. Nevertheless, approximately 20% of electroporation failed due to the absence of transfected cells as revealed by the lack of fluorescent cells (in GFP or mCherry and Smo conditions). These animals were excluded from the study.

### Western blotting

Electroporated zones of the somatosensory cortex were homogenized in RIPA buffer (150 mM NaCl, 1% Triton X-100, 0,1% SDS, 50 mM Tris HCl, pH 8, containing protease inhibitors (Complete Mini; Roche). Lysates were centrifuged (10.000g for 10 min at 4°C) and the supernatant was heated at 90°C for 5 min with Laemmli loading buffer. Loading was 20 µg of proteins as determined using a modified Bradford reaction (BioRad Laboratories). Proteins were separated in 7–15% SDS-PAGE and electrophoretically transferred to nitrocellulose membranes. Membranes were blocked with 5% Bovine Serum Albumin (BSA) in TBS 0,1% Tween 20 (TBST) for 2h at RT, then incubated with primary antibodies diluted in TBST containing 3% BSA overnight at 4°C or 2h at RT. Blots were probed with antibody against KCC2 pSer^940^ (1 mg/ml; rabbit, Novus Biologicals), KCC2 pThr^1007^ (1 mg/ml; sheep, Division of Signal Transduction Therapy Unit (DSTT), University of Dundee), and KCC2 (1:2000; rabbit, US Biological). After washing with TBST, membranes were incubated with HRP-conjugated secondary antibodies diluted in TBST containing 3% BSA for 60 min, washed with TBST and then developed using the G:BOX gel imaging system (Syngene). The appropriate exposure time of the digital camera for acquisition of chemiluminescent signals from immunoblots was adapted to avoid saturated pixels and expression levels were estimated by Image J software (NIH, Bethesda, MD; http://rsb.info.nih.gov/ij/).

### Immunocytochemistry and confocal microscopy

Under deep anesthesia with isoflurane prior to chloral hydrate (7% in PBS 1M), P10 to P30 electroporated rats were intracardially perfused with cold PBS (1M) followed by 4% PFA in PBS. Brains were removed, post-fixed overnight at 4°C and rinsed in PBS. Coronal cortical sections (70μm thick) were obtained using a vibratome (Microm HM 650V). Sections were incubated first for 1 h in PBS with 1% BSA and 0.3% Triton X-100, then overnight at 4°C with rabbit anti-Smo (1:500; ab38686; Abcam) or rabbit anti-caspase-3 cleaved (1:500; 9661S; Cell Signaling) or chicken anti-MAP2 (1:5000; ab5392; Abcam) or mouse anti-NeuN (1:1000; MAB377; Chemicon), or rabbit anti-FoxP2 (1:4000; ab16046; Abcam), or mouse anti-synaptophysin (1:000; MAB5258; Chemicon). Sections were rinsed in PBS and incubated for 2h with the corresponding Alexa488 (1:1000; FluoProbes), Cy5 or Cy3-conjugated secondary antibodies diluted in PBS (1:1000; Chemicon). DAPI staining was applied for nuclear localization (Vector Laboratories H-1200). Control tissues used to determine the level of nonspecific staining included tissues incubated without primary antibody. Sequential acquisition of immunoreactivity of GFP positive pyramidal-like cells was performed using laser scanning confocal microscope (Zeiss LSM 510 Meta) with a 40X or 63X oil-immersion objectives. For each experimental condition, a semi-quantitative analysis of Synaptophysin (Syn) labeled area fraction per field was measured and reported to the area fraction of positive pixels for MAP2. The surface areas of NeuN immuno-positive neurons expressing GFP or Smo-related constructs were measured manually using the Freehands selection tool in Image J. In each set of images, laser light levels and detector gain and offset were adjusted to avoid any saturated levels. Optical sections were digitized (1024 × 1024 pixels) and processed using Image J software. All of the images were analyzed blindly.

### Relative Quantitative Expression of mRNA transcripts

Expressions of Gli1 and Ptch1 mRNA in the electroporated regions of the somatosensory cortex were measured by real-time quantitative RT-PCR. Total RNAs were isolated from cerebral cortices (P15 rats) using Mini RNeasy kit (Qiagen) then converted to cDNA using 1 µg RNA and a QuantiTect Reverse Transcription kit (Qiagen) according to manufacturer’s instructions. PCR was carried out with the LightCycler 480 SYBR Green I Master (Roche Applied Science) with 1 µL cDNA using the following oligonucleotides (QuantiTect Primer Assays, Qiagen): Gli1 (Gli1; QT01290324), Ptch1 (QT01579669), KCC2 (Slc12a5; QT00145327) and glyceraldehyde-3-phosphate dehydrogenase (GAPDH; QT001199633). Relative mRNA values were calculated using the LC480 software and GAPDH as the housekeeping gene. PCR was performed in replicates of 3.

### Sonic hedgehog protein (Shh) immunoassay

Cortical tissues from electroporated rats at the indicated age were homogenized in RIPA buffer (150 mM NaCl, 1% Triton X-100, 0.1% SDS, 50 mM Tris HCl, pH 8, containing protease inhibitors (Complete Mini; Roche)). Lysates were centrifuged (5000 g for 5 min at 4°C). Loading was 200 µg of protein as determined using a modified Bradford reaction (BioRad Laboratories). Quantification of Shh was performed with Rat Shh ELISA Kit (FineTest, Wuhan Fine Biotech Co. Ltd., China) in the concentrated solutions following the manufacturer’s protocol. Experiments and analysis were done blindly.

### Slices preparation and electrophysiological recordings

Electroporated cortical regions of the P14 rat were identified using an atlas of the developing rat brain (Khazipov et al., 2015). Electroporated cortical regions of the P20 and P30 rat were identified using the Paxinos and Watson rat brain atlas (Paxinos and Watson, 2005). Based on these two atlases, the somatosensory regions were prepared for acute slice experiments from P14, P20 and P30 rats in ice-cold (2-4°C) oxygenated modified artificial cerebrospinal fluid (ACSF) (0.5mM CaCl2 and 7mM MgSO4; NaCl replaced by an equimolar concentration of choline). Slices (300 µm thick) were cut with a Vibratome (VT1000E; Leica, Nussloch, Germany) and kept at room temperature (25°C) for at least one hour before recording in oxygenated normal ACSF containing (in mM): 126 NaCl, 3.5 KCl, 2 CaCl_2_, 1.3 MgCl_2_, 1.2 NaH_2_PO_4_, 25 NaHCO_3_ and 11 glucose, pH 7.4 equilibrated with 95% O_2_ and 5% CO_2_. Slices were then transferred to a submerged recording chamber perfused with ACSF (3 mL/min) at 34°C.

### Spiking activity and data analysis

Extracellular field potential recordings were performed from P14, P20 and P30 electroporated somatosensory cortical slices. Extracellular tungsten electrodes of 50µm diameter (California Fine Wire, Grover Beach, CA, USA) were positioned in the V/VI cortical pyramidal cell layer of the transfected area to record the Multi Unit Activity (MUA). Field potentials were recorded using a DAM80 Amplifier (World Precision Instruments, Sarasota, FL, USA) using a 1–3 Hz bandpass filter and analyzed off-line with the Axon package MiniAnalysis program (Jaejin Software, Leonia, NJ, USA). To determine the developmental changes in the GABA_A_ signaling, we used isoguvacine, a potent and selective GABA_A_ receptor agonist (Krogsgaard-Larsen and Johnston, 1978). We determined the effect of isoguvacine on MUA as a ratio of the spiking frequency at the peak of the isoguvacine response to the spiking frequency in control.

### Cl^-^-Sensor fluorescence recordings from brain slices

To perform a non-invasive monitoring of neuronal intracellular chloride concentration ([Cl^-^]_i_) we used a ratiometric genetically-encoded Cl^-^-sensitive probe called Cl-Sensor (Markova et al., 2008; Waseem et al., 2010) that was co-expressed into the cells of interest together with other constructs as described above. The acquisition of fluorescence images was performed using a customized imaging set-up and consecutive cells excitation at 430 and 500 nm and emission at 480 and 540 nm. The frequency of acquisition was 0.05 Hz. The duration of excitation was selected for each cell type and was selected to avoid use-dependent bleaching of the signal (Friedel et al., 2013). Results are expressed as ratio R430/500. Experiments were performed on acute cortical slices from *in utero* electroporated rat pups on postnatal days P10 and P30. Individual slices were transferred to a specially designed recording chamber where they were fully submerged and superfused with oxygenated ACSF complemented with 1 µM tetrodotoxin, 0.3 µM strychnine, and 10 µM NBQX to prevent spontaneous neuronal activity and noncontrolled [Cl^−^]_i_ changes at 30–32°C at a rate of 2–3 ml/min. The applications of the ACSF solution containing isoguvacine (30 µM) or KCl (25 mM) + isoguvacine (30 µM) were performed with a perfusion system.

### Seizure induction with pentylenetetrazol (PTZ)

To evaluate the susceptibility to seizures in controls rats (GFP-transfected brains), in Smo-CA and Smo-DN animals at P30, PTZ (25 mg/kg; Sigma) was administered via intraperitoneal injections every 10 min until generalized seizures occurred. Rats were placed in a plexiglass cage and the time to the onset of the generalized seizure was measured by observation. There was no significant difference in weight and sex between groups of animals. Experiments were done blind and the electroporated region was verified after PTZ induction. Rats with a very focal fluorescence or fluorescence in other brain regions than somatosensory cortex were excluded.

### Primary cultures and transfection of rat hippocampal neurons

Neurons from 18-day-old rat embryos were dissected and dissociated using trypsin and plated at a density of 70,000 cells cm^−2^ in minimal essential medium (MEM) supplemented with 10% NU serum (BD Biosciences, Le Pont de Claix, France), 0.45% glucose, 1 mM sodium pyruvate, 2 mM glutamine and 10 U ml−1 penicillin–streptomycin (Buerli et al., 2007). On days 7, 10 and 13 of culture incubation (DIV, days *in vitro*), half of the medium was changed to MEM with 2% B27 supplement (Invitrogen). For electrophysiology, neuronal cultures were plated on coverslips placed in 35-mm culture dishes. Twelve hours before plating, dishes with coverslips were coated with polyethylenimine (5 mg/ml). Transfection of cultured neurons was performed with 300 µl Opti-MEM medium mixed with 7 µl Lipofectamine 2000 (Invitrogen), 1 µl Magnetofection CombiMag (OZ Biosciences) per µg of DNA and 1.5 µg premixed DNAs encoding constructs of interest. The mixture was incubated for 20 min at room temperature and thereafter distributed dropwise above the neuronal culture. Culture dishes (35-mm) were placed on a magnetic plate (OZ Biosciences) and incubated for 35 min at 37°C, 5% CO_2_. Transfection was terminated by the substitution of 80% of the incubation solution with fresh culture medium. Cells were used in the experiments 48-72 hours after transfection. These experiments were based on co-transfection into the same cell of two different pcDNAs encoding a fluorescent marker of transfection (eGFP or mCherry, 0.3 µg), and Smo-related constructs (1.2 µg).

#### Gramicidin-perforated patch-clamp recordings

Gramicidin-perforated patch-clamp recordings were performed on primary hippocampal neurons, transfected with a mixture of constructs encoding mCherry (mock) or Smo-DN or Smo-CA with or without KCC2-pH_ext_ (Friedel et al., 2017). Measurements were performed 2 or 3 days after transfection (corresponding to 8 or 9 DIV). Coverslips with transfected neurons were placed onto the inverted microscope and perfused with an external Hepes-buffered solution (HBS) (in mM): 140 NaCl, 2.5 KCl, 20 Hepes, 20 d-glucose, 2.0 CaCl_2_, 2.0 MgCl_2_, and 0.02 Bumetanide, pH 7.4. For recording from neurons, external HBS contained 0.5 µM tetrodotoxin and 15 µM bumetanide. The recording micropipettes (5 MΩ) were filled with a solution containing (in mM): 150 KCl, 10 Hepes, and 20 µg/ml gramicidin A, pH 7.2. Isoguvacine (30 µM) was dissolved in an external solution and focally applied to recorded cells through a micropipette connected to a Picospritzer (General Valve Corporation, pressure 5 p.s.i.). Recordings were done using an Axopatch-200A amplifier and pCLAMP acquisition software (Molecular Devices) in voltage-clamp mode. Data were low pass filtered at 2 kHz and acquired at 10 kHz. Isoguvacine responses were recorded at voltages −110, −90, −70 and −50 mV or at −70, −50, −30 and −10 mV depending on neuron GABA inversion potential. A linear regression was used to calculate the best-fit line of the voltage dependence of the isoguvacine responses.

### Surface immunolabeling on living neurons and analysis of KCC2-pH_ext_ proteins

For immunolabeling of KCC2-pH_ext_ proteins on living neurons, rabbit anti-GFP antibody were dissolved in culture media applied to neurons for 2 hours at 37°C, 5% CO2 (Friedel et al., 2015). Neurons were then rinsed three times at room temperature (RT) with HBS, labeled with anti-rabbit Cy3-conjugated antibody for 20 min at 13°C and fixed in Antigenfix (Diapath). To reveal intracellular pool of live-labelled proteins, cells were permeabilized with 0.3% Triton X-100, blocked by 5% goat serum and incubated during 1 h at RT with anti-rabbit Alexa 647-conjugated antibody. For visualization of the total pool of overexpressed KCC2-pH_ext_ cells were labeled overnight (4°C) with mouse anti-GFP antibody and for 1 hour at RT with anti-mouse Alexa 488-conjugated antibody. For control of the cell membrane integrity during live-cell immunolabeling, one batch of cultures were routinely transfected with KCC2-pH_ext_ mutant ΔNTD-KCC2-pH_ext_ that does not incorporate into the plasma membrane (Friedel et al., 2017). KCC2 A/A, a double phosphomimetic mutant of KCC2 for T906A/T1007A that maintains KCC2 at the cell surface (Friedel et al., 2015) was used as a positive control.

Images of labeled neurons were acquired with an Olympus Fluorview-500 confocal microscope (oil-immersion objective 60x (NA1.4); zoom 1-5). We randomly selected and focused on a transfected cell by only visualizing Alexa-488 fluorescence and then acquired Z-stack images of Alexa-488, CY3 and Alexa-647 fluorochromes. Each Z-stack included 10 planes of 1 µm optical thickness and taken at 0.5 µm distance between planes. The cluster properties and fluorescence intensities of each cell were analyzed with Metamorph software. First, we used the logical “NOT” conversion of pairs of Alexa-647 and CY3 images to isolate in each focal plane the Alexa-647 signal that was not overlapping with CY3 fluorescence restricted to plasma membrane. This gave rise to additional images reflecting the fluorescence of the internalized pool of labeled clusters, called thereafter “NOT-conversion”. Second, the arithmetic summation for each Z-stack and channel was performed to collect the whole fluorescence of the different signals (Alexa-488 = total protein fluorescence; CY3 = plasma membrane restricted fluorescence; NOT-conversion = internalized restricted fluorescence; Alexa-647 = all surface labeled fluorescence). Third, a binary mask was created for each cell from Alexa-488 image to isolate the signal coming from the transfected neuron, and the fluorescence parameters (total fluorescence, single cluster fluorescence as well as density and brightness of clusters) were analyzed for each channel (Alexa-488, CY3, NOT-conversion and Alexa-647) in regions overlapping with the binary mask. The analysis parameters were the same for each experiment and all experiments were done blind. After analysis, data were normalized to the mean value of cells transfected with KCC2-pHext + GFP.

### Statistical analysis

No statistical methods were used to predetermine sample sizes. To ensure the consistency and reproducibility of our results, we conducted repeated trials in different cell cultures and acute brain slices prepared from at least three different animals for each experimental condition. If not stated otherwise, statistics are presented as the mean ± the standard deviation (SD) for normally distributed data and as the median only for non-normally distributed data. Experiments with control and Smo electroporated animals were processed at the same time to ensure homogeneity of experimental conditions. Statistical analyses and assessment of normal distribution (Shapiro-Wilk test) were performed with GraphPad Prism (GraphPad software 5.01). For data showing normal distribution, ANOVA and the post hoc Tukey test were used for multiple comparisons between groups, and a paired t-test was used to compare paired data. For data displaying non-normal distribution Kruskal-Wallis test was used to compare three of more independent groups, Mann-Whitney U-test was used for comparison between 2 independent groups and Wilcoxon matched-pairs signed rank test to analyze differences within one group across conditions.

## Acknowledgements

We thank Drs. Y Ben Ari and J.L. Gaiarsa for critical reading of the manuscript; F. Michel at InMAGIC (INMED Imaging Centre) for technical assistance.

## Authors Contributions

QD, IM, YHB and CP designed research. QD, IM, MH, EB, JZ, YHB and CP performed research. QD, YHB and CP analyzed data. QD, IM and CP wrote the paper.

## References

Aguado, F., Carmona, M. A., Pozas, E., Aguiló, A., Martínez-Guijarro, F. J., Alcantara, S., Borrell, V., Yuste, R., Ibañez, C. F. and Soriano, E. (2003). BDNF regulates spontaneous correlated activity at early developmental stages by increasing synaptogenesis and expression of the K+/Cl-co-transporter KCC2. Development 130, 1267–1280.

Al-Ayadhi, L. Y. (2012). Relationship between Sonic hedgehog protein, brain-derived neurotrophic factor and oxidative stress in autism spectrum disorders. Neurochem. Res. 37, 394–400.

Álvarez-Buylla, A. and Ihrie, R. A. (2014). Sonic hedgehog signaling in the postnatal brain. Semin. Cell Dev. Biol. 33, 105–111.

Antonelli, F., Casciati, A. and Pazzaglia, S. (2019). Sonic hedgehog signaling controls dentate gyrus patterning and adult neurogenesis in the hippocampus. Neural Regen Res 14, 59–61.

Araújo, G. L. L., Araújo, J. A. M., Schroeder, T., Tort, A. B. L. and Costa, M. R. (2014). Sonic hedgehog signaling regulates mode of cell division of early cerebral cortex progenitors and increases astrogliogenesis. Front Cell Neurosci 8, 77.

Belgacem, Y. H. and Borodinsky, L. N. (2015). Inversion of Sonic hedgehog action on its canonical pathway by electrical activity. Proc. Natl. Acad. Sci. U.S.A. 112, 4140–4145.

Belgacem, Y. H., Hamilton, A. M., Shim, S., Spencer, K. A. and Borodinsky, L. N. (2016). The Many Hats of Sonic Hedgehog Signaling in Nervous System Development and Disease. J Dev Biol 4,.

Ben-Ari, Y. (2002). Excitatory actions of gaba during development: the nature of the nurture. Nat. Rev. Neurosci. 3, 728–739.

Ben-Ari, Y. and Holmes, G. L. (2005). The multiple facets of gamma-aminobutyric acid dysfunction in epilepsy. Curr. Opin. Neurol. 18, 141–145.

Ben-Ari, Y., Cherubini, E., Corradetti, R. and Gaiarsa, J. L. (1989). Giant synaptic potentials in immature rat CA3 hippocampal neurones. J. Physiol. (Lond.) 416, 303–325.

Ben-Ari, Y., Gaiarsa, J.-L., Tyzio, R. and Khazipov, R. (2007). GABA: a pioneer transmitter that excites immature neurons and generates primitive oscillations. Physiol. Rev. 87, 1215–1284.

Beug, S. T., Parks, R. J., McBride, H. M. and Wallace, V. A. (2011). Processing-dependent trafficking of Sonic hedgehog to the regulated secretory pathway in neurons. Mol. Cell. Neurosci. 46, 583–596.

Bezard, E., Baufreton, J., Owens, G., Crossman, A. R., Dudek, H., Taupignon, A. and Brotchie, J. M. (2003). Sonic hedgehog is a neuromodulator in the adult subthalamic nucleus. FASEB J. 17, 2337–2338.

Bond, C. W., Angeloni, N., Harrington, D., Stupp, S. and Podlasek, C. A. (2013). Sonic Hedgehog regulates brain-derived neurotrophic factor in normal and regenerating cavernous nerves. J Sex Med 10, 730–737.

Bovolenta, R., Zucchini, S., Paradiso, B., Rodi, D., Merigo, F., Navarro Mora, G., Osculati, F., Berto, E., Marconi, P., Marzola, A., et al. (2010). Hippocampal FGF-2 and BDNF overexpression attenuates epileptogenesis-associated neuroinflammation and reduces spontaneous recurrent seizures. J Neuroinflammation 7, 81.

Brambilla, P., Hardan, A., di Nemi, S. U., Perez, J., Soares, J. C. and Barale, F. (2003). Brain anatomy and development in autism: review of structural MRI studies. Brain Res. Bull. 61, 557–569.

Breunig, J. J., Sarkisian, M. R., Arellano, J. I., Morozov, Y. M., Ayoub, A. E., Sojitra, S., Wang, B., Flavell, R. A., Rakic, P. and Town, T. (2008). Primary cilia regulate hippocampal neurogenesis by mediating sonic hedgehog signaling. Proc. Natl. Acad. Sci. U.S.A. 105, 13127–13132.

Briscoe, J. and Thérond, P. P. (2013). The mechanisms of Hedgehog signalling and its roles in development and disease. Nat. Rev. Mol. Cell Biol. 14, 416–429.

Buerli, T., Pellegrino, C., Baer, K., Lardi-Studler, B., Chudotvorova, I., Fritschy, J.-M., Medina, I. and Fuhrer, C. (2007). Efficient transfection of DNA or shRNA vectors into neurons using magnetofection. Nat Protoc 2, 3090–3101.

Chamma, I., Heubl, M., Chevy, Q., Renner, M., Moutkine, I., Eugène, E., Poncer, J. C. and Lévi, S. (2013). Activity-dependent regulation of the K/Cl transporter KCC2 membrane diffusion, clustering, and function in hippocampal neurons. J. Neurosci. 33, 15488–15503.

Charytoniuk, D., Porcel, B., Rodríguez Gomez, J., Faure, H., Ruat, M. and Traiffort, E. (2002). Sonic Hedgehog signalling in the developing and adult brain. J. Physiol. Paris 96, 9–16.

Chen, Y., Sasai, N., Ma, G., Yue, T., Jia, J., Briscoe, J. and Jiang, J. (2011). Sonic Hedgehog dependent phosphorylation by CK1α and GRK2 is required for ciliary accumulation and activation of smoothened. PLoS Biol. 9, e1001083.

Chen, L., Wan, L., Wu, Z., Ren, W., Huang, Y., Qian, B. and Wang, Y. (2017). KCC2 downregulation facilitates epileptic seizures. Sci Rep 7, 156.

Chen, S.-D., Yang, J.-L., Hwang, W.-C. and Yang, D.-I. (2018). Emerging Roles of Sonic Hedgehog in Adult Neurological Diseases: Neurogenesis and Beyond. Int J Mol Sci 19,.

Choi, J., Ababon, M. R., Soliman, M., Lin, Y., Brzustowicz, L. M., Matteson, P. G. and Millonig, H. (2014). Autism associated gene, engrailed2, and flanking gene levels are altered in postmortem cerebellum. PLoS ONE 9, e87208.

Choudhry, Z., Rikani, A. A., Choudhry, A. M., Tariq, S., Zakaria, F., Asghar, M. W., Sarfraz, M. K., Haider, K., Shafiq, A. A. and Mobassarah, N. J. (2014). Sonic hedgehog signalling pathway: a complex network. Ann Neurosci 21, 28–31.

Côme, E., Marques, X., Poncer, J. C. and Lévi, S. (2019). KCC2 membrane diffusion tunes neuronal chloride homeostasis. Neuropharmacology 107571.

de Los Heros, P., Alessi, D. R., Gourlay, R., Campbell, D. G., Deak, M., Macartney, T. J., Kahle, T. and Zhang, J. (2014). The WNK-regulated SPAK/OSR1 kinases directly phosphorylate and inhibit the K+-Cl-co-transporters. Biochem. J. 458, 559–573.

Delmotte, Q., Diabira, D., Belaidouni, Y., Hamze, M., Kochmann, M., Montheil, A., Gaiarsa, J.-L., Porcher, C. and Belgacem, Y. H. (2020). Sonic hedghog signaling agonist (SAG) triggers BDNF secretion and promotes the maturation of GABAergic networks in the postanatal rat hippocampus. Frontiers in Cellular Neuroscience.

Dumon, C., Diabira, D., Chudotvorova, I., Bader, F., Sahin, S., Zhang, J., Porcher, C., Wayman, G., Medina, I. and Gaiarsa, J.-L. (2018). The adipocyte hormone leptin sets the emergence of hippocampal inhibition in mice. Elife 7,.

Falcicchia, C., Paolone, G., Emerich, D. F., Lovisari, F., Bell, W. J., Fradet, T., Wahlberg, L. U. and Simonato, M. (2018). Seizure-Suppressant and Neuroprotective Effects of Encapsulated BDNF-Producing Cells in a Rat Model of Temporal Lobe Epilepsy. Mol Ther Methods Clin Dev 9, 211–224.

Fang, M., Lu, Y., Chen, G.-J., Shen, L., Pan, Y.-M. and Wang, X.-F. (2011). Increased expression of sonic hedgehog in temporal lobe epileptic foci in humans and experimental rats. Neuroscience 182, 62–70.

Feng, S., Ma, S., Jia, C., Su, Y., Yang, S., Zhou, K., Liu, Y., Cheng, J., Lu, D., Fan, L., et al. (2016). Sonic hedgehog is a regulator of extracellular glutamate levels and epilepsy. EMBO Rep. 17, 682–694.

Friedel, P., Bregestovski, P. and Medina, I. (2013). Improved method for efficient imaging of intracellular Cl(−) with Cl-Sensor using conventional fluorescence setup. Front Mol Neurosci 6, 7.

Friedel, P., Kahle, K. T., Zhang, J., Hertz, N., Pisella, L. I., Buhler, E., Schaller, F., Duan, J., Khanna, A. R., Bishop, P. N., et al. (2015). WNK1-regulated inhibitory phosphorylation of the KCC2 cotransporter maintains the depolarizing action of GABA in immature neurons. Sci Signal 8, ra65.

Friedel, P., Ludwig, A., Pellegrino, C., Agez, M., Jawhari, A., Rivera, C. and Medina, I. (2017). A Novel View on the Role of Intracellular Tails in Surface Delivery of the Potassium-Chloride Cotransporter KCC2. eNeuro 4,.

George Paxinos and Watson, C. The Rat Brain in Stereotaxic Coordinates - 7th Edition.

Halepoto, D. M., Bashir, S., Zeina, R. and Al-Ayadhi, L. Y. (2015). Correlation Between Hedgehog (Hh) Protein Family and Brain-Derived Neurotrophic Factor (BDNF) in Autism Spectrum Disorder (ASD). J Coll Physicians Surg Pak 25, 882–885.

Hammond, R., Blaess, S. and Abeliovich, A. (2009). Sonic hedgehog is a chemoattractant for midbrain dopaminergic axons. PLoS ONE 4, e7007.

Harwell, C. C., Parker, P. R. L., Gee, S. M., Okada, A., McConnell, S. K., Kreitzer, A. C. and Kriegstein, A. R. (2012). Sonic hedgehog expression in corticofugal projection neurons directs cortical microcircuit formation. Neuron 73, 1116–1126.

Hildebrand, M. S., Griffin, N. G., Damiano, J. A., Cops, E. J., Burgess, R., Ozturk, E., Jones, N. C., Leventer, R. J., Freeman, J. L., Harvey, A. S., et al. (2016). Mutations of the Sonic Hedgehog Pathway Underlie Hypothalamic Hamartoma with Gelastic Epilepsy. Am. J. Hum. Genet. 99, 423–429.

Hill, S. A., Blaeser, A. S., Coley, A. A., Xie, Y., Shepard, K. A., Harwell, C. C., Gao, W.-J. and Garcia, A. D. R. (2019). Sonic hedgehog signaling in astrocytes mediates cell type-specific synaptic organization. Elife 8,.

Inoue, K., Furukawa, T., Kumada, T., Yamada, J., Wang, T., Inoue, R. and Fukuda, A. (2012). Taurine inhibits K+-Cl-cotransporter KCC2 to regulate embryonic Cl-homeostasis via with-no-lysine (WNK) protein kinase signaling pathway. J. Biol. Chem. 287, 20839–20850.

Jacob, L. and Lum, L. (2007). Deconstructing the hedgehog pathway in development and disease. Science 318, 66–68.

Jeong, J., Mao, J., Tenzen, T., Kottmann, A. H. and McMahon, A. P. (2004). Hedgehog signaling in the neural crest cells regulates the patterning and growth of facial primordia. Genes Dev. 18, 937–951.

Kahle, K. T. and Delpire, E. (2016). Kinase-KCC2 coupling: Clrheostasis, disease susceptibility, therapeutic target. J. Neurophysiol. 115, 8–18.

Kahle, K. T., Deeb, T. Z., Puskarjov, M., Silayeva, L., Liang, B., Kaila, K. and Moss, S. J. (2013). Modulation of neuronal activity by phosphorylation of the K-Cl cotransporter KCC2. Trends Neurosci. 36, 726–737.

Kahle, K. T., Merner, N. D., Friedel, P., Silayeva, L., Liang, B., Khanna, A., Shang, Y., Lachance-Touchette, P., Bourassa, C., Levert, A., et al. (2014). Genetically encoded impairment of neuronal KCC2 cotransporter function in human idiopathic generalized epilepsy. EMBO Rep. 15, 766–774.

Kelsch, W., Hormuzdi, S., Straube, E., Lewen, A., Monyer, H. and Misgeld, U. (2001). Insulin-like growth factor 1 and a cytosolic tyrosine kinase activate chloride outward transport during maturation of hippocampal neurons. J. Neurosci. 21, 8339–8347.

Khazipov, R., Zaynutdinova, D., Ogievetsky, E., Valeeva, G., Mitrukhina, O., Manent, J.-B. and Represa, A. (2015). Atlas of the Postnatal Rat Brain in Stereotaxic Coordinates. Front Neuroanat 9, 161.

Khirug, S., Huttu, K., Ludwig, A., Smirnov, S., Voipio, J., Rivera, C., Kaila, K. and Khiroug, L. (2005). Distinct properties of functional KCC2 expression in immature mouse hippocampal neurons in culture and in acute slices. Eur. J. Neurosci. 21, 899–904.

Kilb, W., Kirischuk, S. and Luhmann, H. J. (2013). Role of tonic GABAergic currents during pre- and early postnatal rodent development. Front Neural Circuits 7, 139.

Kim, J., Kato, M. and Beachy, P. A. (2009). Gli2 trafficking links Hedgehog-dependent activation of Smoothened in the primary cilium to transcriptional activation in the nucleus. Proc. Natl. Acad. Sci. U.S.A. 106, 21666–21671.

Kirmse, K., Kummer, M., Kovalchuk, Y., Witte, O. W., Garaschuk, O. and Holthoff, K. (2015). GABA depolarizes immature neurons and inhibits network activity in the neonatal neocortex in vivo. Nat Commun 6, 7750.

Klioueva, I. A., van Luijtelaar, E. L., Chepurnova, N. E. and Chepurnov, S. A. (2001). PTZ-induced seizures in rats: effects of age and strain. Physiol. Behav. 72, 421–426.

Kriegstein, A. R. and Noctor, S. C. (2004). Patterns of neuronal migration in the embryonic cortex. Trends Neurosci. 27, 392–399.

Krogsgaard-Larsen, P. and Johnston, G. A. (1978). Structure-activity studies on the inhibition of GABA binding to rat brain membranes by muscimol and related compounds. J. Neurochem. 30, 1377–1382.

Kuzirian, M. S. and Paradis, S. (2011). Emerging themes in GABAergic synapse development. Prog. Neurobiol. 95, 68–87.

Lee, H. H. C., Walker, J. A., Williams, J. R., Goodier, R. J., Payne, J. A. and Moss, S. J. (2007). Direct protein kinase C-dependent phosphorylation regulates the cell surface stability and activity of the potassium chloride cotransporter KCC2. J. Biol. Chem. 282, 29777–29784.

Lee, H. H. C., Deeb, T. Z., Walker, J. A., Davies, P. A. and Moss, S. J. (2011). NMDA receptor activity downregulates KCC2 resulting in depolarizing GABAA receptor-mediated currents. Nat. Neurosci. 14, 736–743.

Leonzino, M., Busnelli, M., Antonucci, F., Verderio, C., Mazzanti, M. and Chini, B. (2016). The Timing of the Excitatory-to-Inhibitory GABA Switch Is Regulated by the Oxytocin Receptor via KCC2. Cell Rep 15, 96–103.

Logue, S. E. and Martin, S. J. (2008). Caspase activation cascades in apoptosis. Biochem. Soc. Trans. 36, 1–9.

Ludwig, A., Uvarov, P., Soni, S., Thomas-Crusells, J., Airaksinen, M. S. and Rivera, C. (2011). Early growth response 4 mediates BDNF induction of potassium chloride cotransporter 2 transcription. J. Neurosci. 31, 644–649.

Lüttjohann, A., Fabene, P. F. and van Luijtelaar, G. (2009). A revised Racine’s scale for PTZ-induced seizures in rats. Physiol. Behav. 98, 579–586.

Maisonpierre, P. C., Belluscio, L., Friedman, B., Alderson, R. F., Wiegand, S. J., Furth, M. E., Lindsay, R. M. and Yancopoulos, G. D. (1990). NT-3, BDNF, and NGF in the developing rat nervous system: parallel as well as reciprocal patterns of expression. Neuron 5, 501–509.

Markova, O., Mukhtarov, M., Real, E., Jacob, Y. and Bregestovski, P. (2008). Genetically encoded chloride indicator with improved sensitivity. J. Neurosci. Methods 170, 67–76.

Medina, I., Friedel, P., Rivera, C., Kahle, K. T., Kourdougli, N., Uvarov, P. and Pellegrino, C. (2014). Current view on the functional regulation of the neuronal K(+)-Cl(−) cotransporter KCC2. Front Cell Neurosci 8, 27.

Memi, F., Zecevic, N. and Radonjić, N. (2018). Multiple roles of Sonic Hedgehog in the developing human cortex are suggested by its widespread distribution. Brain Struct Funct 223, 2361–2375.

Menshanov, P. N., Lanshakov, D. A. and Dygalo, N. N. (2015). proBDNF is a major product of bdnf gene expressed in the perinatal rat cortex. Physiol Res 64, 925–934.

Mitchell, N., Petralia, R. S., Currier, D. G., Wang, Y.-X., Kim, A., Mattson, M. P. and Yao, P. J. (2012). Sonic hedgehog regulates presynaptic terminal size, ultrastructure and function in hippocampal neurons. J. Cell. Sci. 125, 4207–4213.

Moore, Y. E., Kelley, M. R., Brandon, N. J., Deeb, T. Z. and Moss, S. J. (2017). Seizing Control of KCC2: A New Therapeutic Target for Epilepsy. Trends Neurosci. 40, 555–571.

Moore, Y. E., Deeb, T. Z., Chadchankar, H., Brandon, N. J. and Moss, S. J. (2018). Potentiating KCC2 activity is sufficient to limit the onset and severity of seizures. Proc. Natl. Acad. Sci. U.S.A. 115, 10166–10171.

Mueller, T. M., Remedies, C. E., Haroutunian, V. and Meador-Woodruff, J. H. (2015). Abnormal subcellular localization of GABAA receptor subunits in schizophrenia brain. Transl Psychiatry 5, e612.

Paradiso, B., Marconi, P., Zucchini, S., Berto, E., Binaschi, A., Bozac, A., Buzzi, A., Mazzuferi, M., Magri, E., Navarro Mora, G., et al. (2009). Localized delivery of fibroblast growth factor-2 and brain-derived neurotrophic factor reduces spontaneous seizures in an epilepsy model. Proc. Natl. Acad. Sci. U.S.A. 106, 7191–7196.

Parra, L. M. and Zou, Y. (2010). Sonic hedgehog induces response of commissural axons to Semaphorin repulsion during midline crossing. Nat. Neurosci. 13, 29–35.

Pascual, O., Traiffort, E., Baker, D. P., Galdes, A., Ruat, M. and Champagnat, J. (2005). Sonic hedgehog signalling in neurons of adult ventrolateral nucleus tractus solitarius. Eur. J. Neurosci. 22, 389–396.

Petralia, R. S., Schwartz, C. M., Wang, Y.-X., Mattson, M. P. and Yao, P. J. (2011). Subcellular localization of Patched and Smoothened, the receptors for Sonic hedgehog signaling, in the hippocampal neuron. J. Comp. Neurol. 519, 3684–3699.

Petralia, R. S., Wang, Y.-X., Mattson, M. P. and Yao, P. J. (2012). Subcellular distribution of patched and smoothened in the cerebellar neurons. Cerebellum 11, 972–981.

Qin, S., Sun, D., Zhang, C., Tang, Y., Zhou, F., Zheng, K., Tang, R. and Zheng, Y. (2019). Downregulation of sonic hedgehog signaling in the hippocampus leads to neuronal apoptosis in high-fat diet-fed mice. Behav. Brain Res. 367, 91–100.

Radzikinas, K., Aven, L., Jiang, Z., Tran, T., Paez-Cortez, J., Boppidi, K., Lu, J., Fine, A. and Ai, X. (2011). A Shh/miR-206/BDNF cascade coordinates innervation and formation of airway smooth muscle. J. Neurosci. 31, 15407–15415.

Riffault, B., Kourdougli, N., Dumon, C., Ferrand, N., Buhler, E., Schaller, F., Chambon, C., Rivera, C., Gaiarsa, J.-L. and Porcher, C. (2018). Pro-Brain-Derived Neurotrophic Factor (proBDNF)-Mediated p75NTR Activation Promotes Depolarizing Actions of GABA and Increases Susceptibility to Epileptic Seizures. Cereb. Cortex 28, 510–527.

Riobo, N. A., Saucy, B., Dilizio, C. and Manning, D. R. (2006). Activation of heterotrimeric G proteins by Smoothened. Proc. Natl. Acad. Sci. U.S.A. 103, 12607–12612.

Rivell, A., Petralia, R. S., Wang, Y.-X., Clawson, E., Moehl, K., Mattson, M. P. and Yao, P. J. (2019). Sonic hedgehog expression in the postnatal brain. Biol Open 8,.

Rivera, C., Voipio, J., Payne, J. A., Ruusuvuori, E., Lahtinen, H., Lamsa, K., Pirvola, U., Saarma, M. and Kaila, K. (1999). The K+/Cl-co-transporter KCC2 renders GABA hyperpolarizing during neuronal maturation. Nature 397, 251–255.

Ruat, M., Roudaut, H., Ferent, J. and Traiffort, E. (2012). Hedgehog trafficking, cilia and brain functions. Differentiation 83, S97–104.

Saitsu, H., Sonoda, M., Higashijima, T., Shirozu, H., Masuda, H., Tohyama, J., Kato, M., Nakashima, M., Tsurusaki, Y., Mizuguchi, T., et al. (2016). Somatic mutations in GLI3 and OFD1 involved in sonic hedgehog signaling cause hypothalamic hamartoma. Ann Clin Transl Neurol 3, 356–365.

Sawano, E., Takahashi, M., Negishi, T. and Tashiro, T. (2013). Thyroid hormone-dependent development of the GABAergic pre- and post-synaptic components in the rat hippocampus. Int. J. Dev. Neurosci. 31, 751–761.

Sernagor, E., Chabrol, F., Bony, G. and Cancedda, L. (2010). GABAergic control of neurite outgrowth and remodeling during development and adult neurogenesis: general rules and differences in diverse systems. Front Cell Neurosci 4, 11.

Silayeva, L., Deeb, T. Z., Hines, R. M., Kelley, M. R., Munoz, M. B., Lee, H. H. C., Brandon, N. J., Dunlop, J., Maguire, J., Davies, P. A., et al. (2015). KCC2 activity is critical in limiting the onset and severity of status epilepticus. Proc. Natl. Acad. Sci. U.S.A. 112, 3523–3528.

Spoljaric, A., Seja, P., Spoljaric, I., Virtanen, M. A., Lindfors, J., Uvarov, P., Summanen, M., Crow, A. K., Hsueh, B., Puskarjov, M., et al. (2017). Vasopressin excites interneurons to suppress hippocampal network activity across a broad span of brain maturity at birth. Proc. Natl. Acad. Sci. U.S.A. 114, E10819–E10828.

Su, Y., Yuan, Y., Feng, S., Ma, S. and Wang, Y. (2017). High frequency stimulation induces sonic hedgehog release from hippocampal neurons. Sci Rep 7, 43865.

Traiffort, E., Angot, E. and Ruat, M. (2010). Sonic Hedgehog signaling in the mammalian brain. J. Neurochem. 113, 576–590.

Tyzio, R., Cossart, R., Khalilov, I., Minlebaev, M., Hübner, C. A., Represa, A., Ben-Ari, Y. and Khazipov, R. (2006). Maternal oxytocin triggers a transient inhibitory switch in GABA signaling in the fetal brain during delivery. Science 314, 1788–1792.

Ung, D. C., Iacono, G., Méziane, H., Blanchard, E., Papon, M.-A., Selten, M., van Rhijn, J.-R., Montjean, R., Rucci, J., Martin, S., et al. (2018). Ptchd1 deficiency induces excitatory synaptic and cognitive dysfunctions in mouse. Mol. Psychiatry 23, 1356–1367.

Vernadakis, A. and Woodbury, D. M. (1969). The developing animal as a model. Epilepsia 10, 163–178.

Wang, D. D. and Kriegstein, A. R. (2009). Defining the role of GABA in cortical development. J. Physiol. (Lond.) 587, 1873–1879.

Wang, B., Zhang, Y., Dong, H., Gong, S., Wei, B., Luo, M., Wang, H., Wu, X., Liu, W., Xu, X., et al. (2018). Loss of Tctn3 causes neuronal apoptosis and neural tube defects in mice. Cell Death Dis 9, 520.

Waseem, T., Mukhtarov, M., Buldakova, S., Medina, I. and Bregestovski, P. (2010). Genetically encoded Cl-Sensor as a tool for monitoring of Cl-dependent processes in small neuronal compartments. J. Neurosci. Methods 193, 14–23.

Wu, C. and Sun, D. (2015). GABA receptors in brain development, function, and injury. Metab Brain Dis 30, 367–379.

Yamada, J., Okabe, A., Toyoda, H., Kilb, W., Luhmann, H. J. and Fukuda, A. (2004). Cluptake promoting depolarizing GABA actions in immature rat neocortical neurones is mediated by NKCC1. J. Physiol. (Lond.) 557, 829–841.

Yao, P. J., Petralia, R. S., Ott, C., Wang, Y.-X., Lippincott-Schwartz, J. and Mattson, M. P. (2015). Dendrosomatic Sonic Hedgehog Signaling in Hippocampal Neurons Regulates Axon Elongation. J. Neurosci. 35, 16126–16141.

Yao, P. J., Petralia, R. S. and Mattson, M. P. (2016). Sonic Hedgehog Signaling and Hippocampal Neuroplasticity. Trends Neurosci. 39, 840–850.

Zhang, X. M., Ramalho-Santos, M. and McMahon, A. P. (2001). Smoothened mutants reveal redundant roles for Shh and Ihh signaling including regulation of L/R asymmetry by the mouse node. Cell 105, 781–792.

Zuñiga, N. R. and Stoeckli, E. T. (2017). Sonic -’Jack-of-All-Trades’ in Neural Circuit Formation. J Dev Biol 5,.

